# Impact of 2′-deoxyribo-purine substrates on nonenzymatic RNA template-directed primer extension

**DOI:** 10.1101/2025.08.29.673048

**Authors:** Ziyuan Fang, Orhan Acikgoz, Xiwen Jia, Jahmyl Essex, Ruby Wen, Jack W. Szostak

**Author notes:** Ziyuan Fang, Quantum-Si, Branford, CT, 06405, USA. Xiwen Jia, Novo Nordisk, Lexington, MA, 02421, USA.

## Abstract

The composition of the primordial genetic material remains uncertain. Studies of duplex structure and stability, and of nonenzymatic template copying chemistry, provide insight into the viability of potentially primordial genetic polymers. Recent work suggests that 2′- deoxyribo-purine nucleotides may have been generated together with ribonucleotides on the early Earth. Since DNA/RNA duplexes are known to be less stable than RNA/RNA duplexes, we have examined the impact of dA, dI, and dG substitutions on RNA structure and nonenzymatic template copying. We find that single 2′-deoxyribo-purine substitutions reduce RNA duplex stability, as expected. Crystallographic studies show that such substitutions lead to minimal structural changes but point to diminished solvation as a likely reason for duplex destabilization. Kinetic studies show that dI and dG substrates exhibit slightly weaker template binding and slower rates of template-directed primer extension than the corresponding ribo-purine substrates. In contrast, dA substrates exhibit much slower reaction kinetics but higher template affinity than rA substrates. Our results suggest that a mixed RNA/DNA primordial genetic polymer would have suffered from moderately slower rates of template copying, but that this could have been offset by an advantage due to more facile strand separation or exchange.

**GRAPHICAL ABSTRACT:** 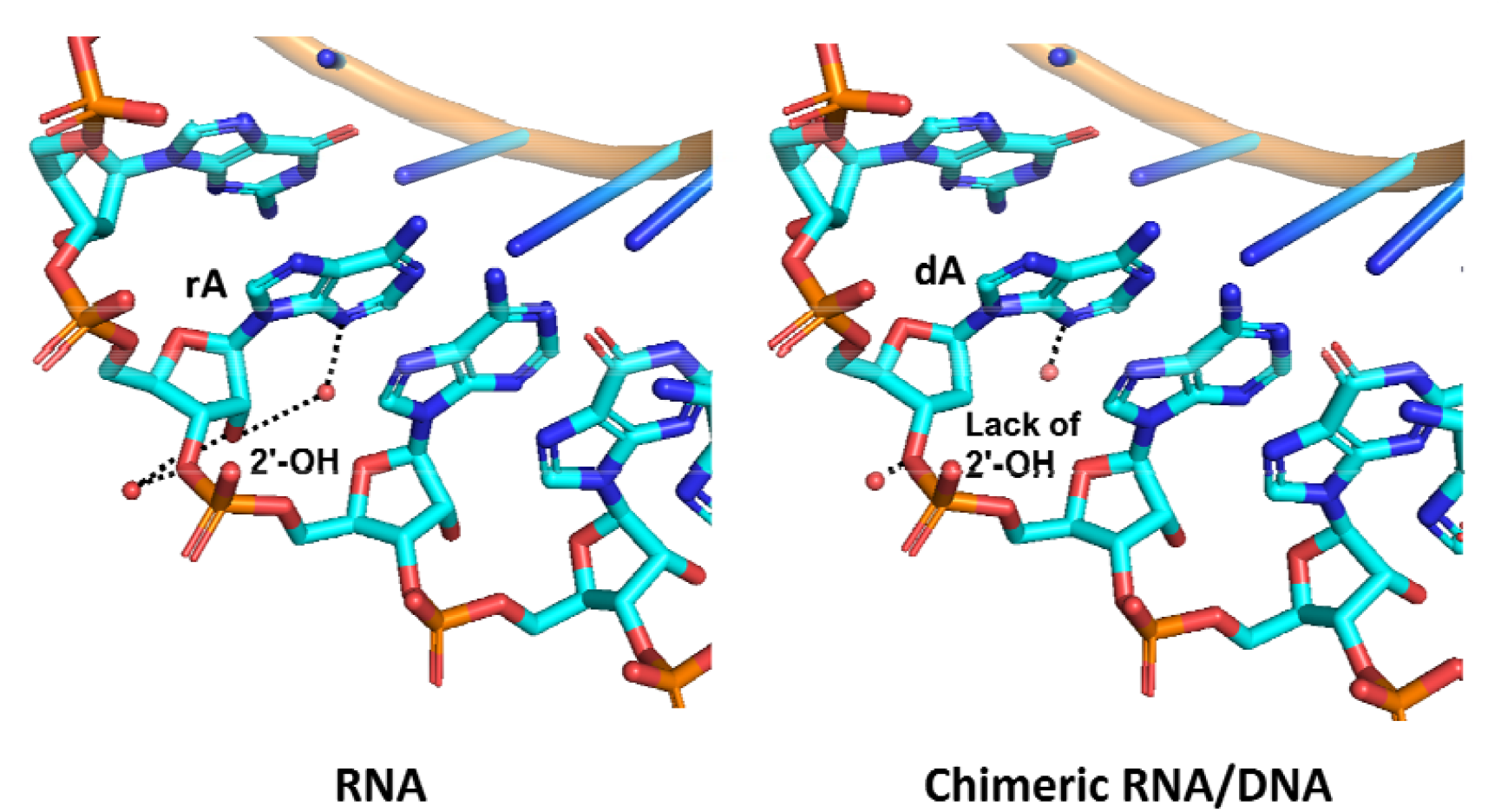

## INTRODUCTION

The RNA World hypothesis proposes that life on Earth originated with RNA serving as both the genetic and functional material of primitive protocells (1,2). However, given the likely chemical complexity of the prebiotic environment, it is possible that 2′-deoxyribo- (3,4), threo- (5-7), arabino- (8-10) and other nucleotides were incorporated into the primordial genetic material alongside ribonucleotides (11-13). Among these candidates, the 2′- deoxyribonucleotides stand out due to the role of DNA in modern biology. The “RNA/DNA World” hypothesis posits that life originated not from pure RNA but from a chimeric system composed of both RNA and DNA (4,14,15). This hypothesis is an attractive alternative to the traditional genetic takeover hypothesis, in which DNA gradually supplanted RNA as the genetic material while RNA retained catalytic functions until the eventual rise of proteins (16,17). Instead, the RNA/DNA model suggests that RNA and DNA diverged from a shared ancestral polymer and evolved toward distinct roles in information storage and catalytic functionality (18). Recent experimental findings support the possibility of prebiotic synthetic routes to 2′- deoxyadenosine (dA) and 2′-deoxyinosine (dI) (19-21). Thus at least some DNA building blocks may have been available before the advent of the complex enzymes that catalyze the biological conversion of ribo- to 2′-deoxyribonucleotides (22,23). This possibility raises the question of whether the incorporation of some 2′-deoxyribonucleotides into primordial RNA would have facilitated or inhibited primordial nucleic acid replication.

The absence of a 2′-hydroxyl group in deoxyribonucleotides renders DNA considerably less labile to strand cleavage than RNA under a range of conditions (24). This stability is especially relevant in the context of nonenzymatic RNA replication, which requires high concentrations of divalent metal ions such as Mg^2+^ to catalyze primer extension (25). However, divalent metal ions also catalyze RNA cleavage by transesterification (26). Consequently, the incorporation of 2′-deoxyribonucleotides into early nucleic acid systems may have helped to preserve the integrity of longer oligonucleotides. This stability may have been critical for the formation and persistence of functional sequences such as ribozymes. Furthermore, chimeric RNA/DNA strands are known to exhibit significantly reduced duplex thermal stability, with this destabilization being particularly pronounced when all purine ribonucleotides are replaced by their deoxyribonucleotide counterparts (4). This duplex destabilization could potentially facilitate the strand separation and exchange processes that are essential for the virtual circular genome (VCG) mode of primordial replication (27,28).

Another interesting possibility arises from the fact that purine substrates exhibit substantially higher template affinities compared to pyrimidine substrates, likely due to the stronger base-stacking interactions of purines (29). This difference could lead to biased or error-prone template copying. For example, the predominant errors in nonenzymatic template copying result from the incorporation of G opposite A or U in the template. We have therefore asked whether replacing the purine ribonucleotides with 2′-deoxyribonucleotides would reduce this difference in substrate affinities, potentially leading to more uniform copying kinetics.

In this study, we have investigated the effects of 2′-deoxyribo-purines on nonenzymatic template copying by conducting thermodynamic, structural, and kinetic analyses. We find that the incorporation of a single 2′-deoxyribo-purine nucleotide reduces the thermodynamic stability of RNA duplexes. Structural analyses of RNA duplexes containing 2′- deoxyribo-purine nucleotides suggest that the observed destabilization is likely due to reduced solvation effects, since base pairing and sugar conformations are unchanged. However, our kinetic studies of nonenzymatic primer extension show that the incorporation of 2′-deoxyribo-purines does not reduce the large difference in binding affinities between purine and pyrimidine bridged-dinucleotide substrates. Our results suggest that a primordial genetic polymer containing both ribo- and 2′-deoxyribo-nucleotides would likely behave similarly to RNA but with somewhat slower template copying chemistry, offset by a possible advantage in strand separation and exchange, which could in turn facilitate cycles of replication.

## MATERIAL AND METHODS

### General Information

All chemicals were purchased from Sigma-Aldrich (St. Louis, MO) and used without purification unless otherwise noted. Phosphoramidites and reagents used for solid-phase RNA synthesis were purchased from ChemGenes (Wilmington, MA) and Glen Research (Sterling, MA). Preparatory-scale high-performance liquid chromatography (HPLC) was carried out on an Agilent 1290 HPLC system, equipped with a preparative-scale Agilent ZORBAX Eclipse-XDB C18 column (21.2 × 250mm, 7 µm particle size). Purity of synthesized products was determined either by NMR or high-resolution mass spectrometry (HRMS). ^1^H and ^31^P spectra (400 MHz for ^1^H, 162 MHz for ^31^P) were acquired on a Bruker Ascend 9.4 T/400 MHz NMR spectrometer equipped with a Bruker SampleCase Plus autosampler at 25 °C. HRMS was carried out on an Agilent 6520 QTOF LC-MS.

### Oligonucleotide Synthesis

Oligonucleotides were synthesized with DMT-off on a K&A H-8-SE-Oligo Synthesizer, then cleaved from the solid support and deprotected with ammonium hydroxide solution at room temperature overnight. The mixtures were lyophilized and then incubated with triethylamine trihydrofluoride (room temperature 3 days for s^2^C and s^2^U-containing oligonucleotides, 65 °C 2.5 h for canonical oligonucleotides) to remove the 2′-TBDMS protecting group. The oligonucleotides were then purified by PAGE. The purity of oligonucleotides was confirmed by LC-MS on an Agilent 6520 TOF mass spectrometer.

### Melting Temperatures of RNA Duplexes

Melting temperatures were measured using an Agilent Cary 3500 UV-Vis Spectrophotometer. For each pair of complementary oligonucleotides, 250 μL samples were prepared with the desired concentration of oligonucleotide in 10 mM Tris-HCl (pH 8.0), 1 M NaCl, and 2.5 mM EDTA. 200 μL mineral oil was added to the top of the RNA solution in the cuvette (Starna 16.100-Q-10/Z15-GL14-C) to prevent the evaporation of water. Melting curves were collected by following absorbance at 260 nm as a function of temperature using a temperature ramp of 0.2 °C/min. The readings were collected in heating–cooling cycles with respect to a control sample containing 10 mM Tris-HCl (pH 8.0), 1 M NaCl, and 2.5 mM EDTA. The melting temperatures were calculated from the interpolation of sigmoidal curves. For each concentration, two samples were prepared, and for each sample two up and down ramp cycles were carried out, generating 8 datasets per condition, i.e. four datasets from low to high temperature and four datasets from high to low temperature.

### Crystallization

*Crystallization of 16-mer self-complementary RNA duplexes*. 0.33 mM self-complementary 16-mer RNA sequences in nuclease-free water (Invitrogen, Waltham, MA) were heated up to 90 °C for 2 min and then slowly cooled to room temperature. Crystal Screen HT, Index HT, Natrix HT (Hampton Research, Aliso Viejo, CA) and Nuc-Pro HTS (Jena Bioscience, Jena, Germany) kits were used to screen crystallization conditions at 20 °C using the sitting-drop vapor diffusion method. An NT8 robotic system and Rock Imager (Formulatrix, Waltham, MA) were used for crystallization screening and for monitoring the crystal growth process. The oligonucleotide sequences and the optimal crystallization conditions are listed in Supporting Information Tables S1 and S2.

*Crystal Soaking with ribo- or 2′*-*deoxyribo-guanosine-5′-phosphoro-2-aminoimidazolide*. 1 mM self-complementary 14-mer RNA sequences including 4 LNA nucleotides on the 5′ end and two nucleotide 5′-overhangs were mixed with an equal volume of 50 mM dGMP and heated to 90°C for 2 min before being slowly cooled to room temperature. Optimal crystals grew in a crystallization buffer of 10% w/v polyethylene glycol 6,000, 50 mM HEPES pH 7.0, 200 mM ammonium acetate and 150 mM magnesium acetate. To soak 2-AIprG or 2-AIpdG into crystals of the RNA duplex with dGMP, the crystals were transferred to a drop containing crystallization buffer and 25 mM 2-AIprG or 2-AIpdG and incubated for 2-4 hours to allow diffusion of the molecules into the crystal. After incubation with the activated substrates, the crystal was placed in liquid nitrogen.

### X-Ray Diffraction Data Collection, Structure Determination, and Refinement

Diffraction data were collected at a wavelength of ∼1 Å (detailed information is listed in the SI Appendix) under a cold nitrogen gas stream at 99 K on Beamline 501, 503 or 821 at the Advanced Light Source in the Lawrence Berkeley National Laboratory, beamline 17ID-2 at National Synchrotron Light Source II (NSLS-II) and beamline 23ID-B at the Advanced Photon Source (APS). The crystals were exposed for 0.25 s per image with a 0.25 Å oscillation angle. The distances between detector and the crystal were set to 180 to 300 mm. The data were processed by HKL2000 (30) or XDS (31). The structures were solved by molecular replacement using PHASER (32) with the structure of 3ND4 as the search model (33). All structures were refined by Phenix (34) and/or Refmac in CCP4i (35). After several cycles of refinement, water molecules and metal atoms with well-defined density were added in Coot (36). Data collection, phasing, and refinement statistics of the determined structures are listed in Supporting Information Tables S3 and S4.

### Synthesis, purification, and characterization of 5′-5′ imidazolium-bridged dinucleotides (N*N)

The synthesis and purification of 2-aminoimidazolium-bridged dinucleotides (A*A, dA*dA U*U, G*G, dG*dG, C*C, I*I, dI*dI, s^2^C*s^2^C, and s^2^U*s^2^U) were carried out as previously described (29). These bridged dinucleotides were characterized by NMR and HRMS (see Supporting Information).

### Nonenzymatic Primer Extension Reactions

Annealing mixtures containing primer/template/blocker complexes were prepared at 5X final concentration: 7.5 μM 5′-FAM labelled primer, 12.5 μM template, 17.5 μM blocker, 50 mM Tris-Cl pH 8.0, 50 mM NaCl, and 1 mM EDTA. The solution was heated to 85 °C for 30 s and then gradually cooled to 25 °C at a rate of 0.1 °C per second using a thermal cycler. This annealed mixture was then diluted fivefold with a buffer containing 250 mM Tris-Cl pH 8.0, and 125 mM MgCl_2_ to achieve final concentrations of 1.5 μM primer, 2.5 μM template, 3.5 μM blocker, 210 mM Tris-Cl pH 8.0, and 100 mM MgCl_2_. Freshly prepared stock solutions of bridged dinucleotides at 2X desired final concentrations were added to the annealed primer/template/blocker solution to initiate templated primer extension reactions. At each time point, a 0.5 μL aliquot was added to 25 μL of quenching buffer, which contained 25 mM EDTA, 1X TBE, and 4 μM of a DNA sequence complementary to the template, in formamide. Primer extension products were resolved by 20% (19:1) denaturing PAGE. The gel was scanned using an Amersham Typhoon Biomolecular Imager, and the bands were quantified using ImageQuant TL software. Oligonucleotide sequences are provided in Supporting Information Tables S5 and S6.

## RESULTS

### Thermodynamic Analysis of RNA Duplexes containing 2′-deoxyribo-purine Nucleotides

We measured the melting temperatures (*T*_m_) of 9-bp RNA duplexes containing a variable central base pair flanked by constant sequences so that we could evaluate the energetics of 2′-deoxyribo-purine containing base pairs and compare them with the ribo-purine containing base pairs in the same context. *T*_m_ values were measured by variable temperature UV absorbance in 10 mM Tris-HCl at pH 8.0, 1 M NaCl, and 2.5 mM EDTA, at a series of concentrations ranging from 1.25 to 20 μM total RNA (Figure S1). We evaluated the thermodynamic parameters Δ*H*°, Δ*S*°, and Δ*G*° by fitting the melting temperatures at different oligonucleotide concentrations to the Van′t Hoff equation (Figure S2). The resulting thermodynamic data for duplexes with different central base pairs are presented in Table 1. Our previous thermodynamic data with ribo-purine containing base pairs are also listed in Table 1 for comparison (37,38).

**Table 1.** Thermodynamic Parameters of RNA Duplex Formation by Thermal Denaturation

| Entry | Base pair | Sequence | $T_m^a$ (°C) | $\Delta H^{\circ b}$<br>(kcal mol<sup>-1</sup>) | $\Delta S^{\circ b}$<br>(kcal K<sup>-1</sup> mol<sup>-1</sup>) | $\Delta G^{\circ}_{25^\circ C^c}$<br>(kcal mol<sup>-1</sup>) | $\Delta G^{\circ}_{65^\circ C^c}$<br>(kcal mol<sup>-1</sup>) |
| --- | --- | --- | --- | --- | --- | --- | --- |
| **1** | A:U<sup>d</sup> | 5'-CUGA <u>A</u> GUAG-3'<br>3'-GACU <u>U</u> CAUC-5' | 46.8(2) | -80.9(1.0) | -0.226(3) | -13.6(2) | -4.5(1) |
| **2** | dA:U | 5'-CUGA d<u>A</u> GUAG-3'<br>3'-GACU <u>U</u> CAUC-5' | 42.8(1) | -77.5(1.0) | -0.218(3) | -12.4(1) | -3.68(4) |
| **3** | G:C<sup>d</sup> | 5'-CUGA <u>G</u> GUAG-3'<br>3'-GACU <u>C</u> CAUC-5' | 55.6(2) | -76.2(1.3) | -0.205(4) | -15.2(3) | -7.0(1) |
| **4** | dG:C | 5'-CUGA d<u>G</u> GUAG-3'<br>3'-GACU <u>C</u> CAUC-5' | 51.7(1) | -80.1(1.1) | -0.219(3) | -14.6(2) | -5.9(1) |
| **5** | I:C<sup>d</sup> | 5'-CUGA <u>I</u> GUAG-3'<br>3'-GACU <u>C</u> CAUC-5' | 46.2(2) | -74.9(1.2) | -0.208(3) | -13.0(2) | -4.7(1) |
| **6** | dI:C | 5'-CUGA d<u>I</u> GUAG-3'<br>3'-GACU <u>C</u> CAUC-5' | 42.0(1) | -76.9(1.1) | -0.217(3) | -12.2(2) | -3.52(5) |
| **7** | A:s<sup>2</sup>U<sup>d</sup> | 5'-CUGA A GUAG-3'<br>3'-GACU s<sup>2</sup>U CAUC-5' | 53.1(2) | -77.7(1.0) | -0.211(3) | -14.7(2) | -6.3(1) |
| **8** | dA:s<sup>2</sup>U | 5'-CUGA dA GUAG-3'<br>3'-GACU s<sup>2</sup>U CAUC-5' | 50.7(2) | -71.5(0.8) | -0.195(2) | -13.3(1) | -5.5(1) |
| **9** | G:s<sup>2</sup>C<sup>d</sup> | 5'-CUGA G GUAG-3'<br>3'-GACU s<sup>2</sup>C CAUC-5' | 50.1(2) | -72.8(1.0) | -0.198(3) | -13.7(2) | -5.8(1) |
| **10** | dG:s<sup>2</sup>C | 5'-CUGA dG GUAG-3'<br>3'-GACU s<sup>2</sup>C CAUC-5' | 46.1(1) | -77.6(1.0) | -0.216(3) | -13.2(2) | -4.6(1) |
| **11** | I:s<sup>2</sup>C<sup>d</sup> | 5'-CUGA <u>I</u> GUAG-3'<br>3'-GACU s<sup>2</sup>C CAUC-5' | 52.0(2) | -79.0(1.4) | -0.216(4) | -14.6(3) | -6.0(1) |
| **12** | dI:s<sup>2</sup>C | 5'-CUGA d<u>I</u> GUAG-3'<br>3'-GACU s<sup>2</sup>C CAUC-5' | 48.3(1) | -73.9(1.0) | -0.203(2) | -13.4(2) | -5.3(1) |
| **13** | G:U | 5'-CUGA G GUAG-3'<br>3'-GACU U CAUC-5' | 43.2(1) | -82.3(1.0) | -0.233(3) | -12.8(2) | -3.46(4) |
| **14** | dG:U | 5'-CUGA dG GUAG-3'<br>3'-GACU U CAUC-5' | 39.4(1) | -87.4(1.2) | -0.253(4) | -12.1(2) | -1.99(3) |
| **15** | s<sup>2</sup>U:s<sup>2</sup>U<sup>e</sup> | 5'-CUGA s<sup>2</sup>U GUAG-3'<br>3'-GACU s<sup>2</sup>U CAUC-5' | 45.4(2) | -73.4(3.5) | -0.203(11) | -12.7(2) | -4.6(2) |

<sup>a</sup>The reported $T_m$ was calculated from sigmoidal curves of raw thermal UV-vis data at 5 μM total oligonucleotide, 10 mM Tris-HCl 8.0, 1 M NaCl, and 2.5 mM EDTA (Supporting Figure S1).
<sup>b</sup> $\Delta H^\circ$ and $\Delta S^\circ$ were derived from linear fits of Van't Hoff plots of $T_m^{-1}$ versus $\ln(C_T/4)$ , where $C_T$ is the total oligonucleotide concentration (Supporting Figure S2).
<sup>c</sup> $\Delta G^{\circ}_{25^{\circ}\text{C}}$ and $\Delta G^{\circ}_{65^{\circ}\text{C}}$ was calculated from $\Delta H^{\circ}$ and $\Delta S^{\circ}$ according to the equation $\Delta G^{\circ} = \Delta H^{\circ} - T\Delta S^{\circ}$ , where $T = 298.15\text{ K}$ or $338.15\text{ K}$ . Standard errors ( $N = 8$ ) are reported.
<sup>d</sup>The values for G:C, A:U, I:C, G:s<sup>2</sup>C, A:s<sup>2</sup>U, I:s<sup>2</sup>C are reproduced from Table 1 in ref (37) with permission under a Creative Commons Attribution 4.0 International License. Copyright 2025 National Academy of Sciences.
<sup>e</sup>The values for s<sup>2</sup>U:s<sup>2</sup>U are reproduced from Table 1 in ref (38) with permission under a Creative Commons Attribution 4.0 International License. Copyright 2024 American Chemical Society.

Notably, all duplexes containing base pairs with 2′-deoxyribo-purines are weaker than their ribonucleotide counterparts, including both the canonical and 2-thiopyrimidine-modified base pairs, with a relative difference in Δ*G*°_25°C_ of approximately 0.5-1.4 kcal/mol. However, different 2′-deoxyribo-purines reduce base pair stabilities in distinct ways. Base pairs involving deoxyadenosine (dA) and deoxyinosine (dI), especially those paired with 2-thiopyrimidines (dA:s^2^U and dI:s^2^C), exhibit enthalpic penalties accompanied by entropic gains, relative to their ribonucleotide counterparts (A:s^2^U and I:s^2^C). The net effect is a Δ*G*° _25°C_ penalty of approximately 1.2–1.4 kcal/mol. Due to their similar thermodynamic trends and compensatory changes, dA:s^2^U and dI:s^2^C, like A:s^2^U and I:s^2^C, are essentially isoenergetic. In contrast, base pairs involving deoxyguanosine (dG) display enthalpic gains but entropic penalties compared to their G-containing counterparts. The net Δ*G*°_25°C_ penalty for base pairs with dG is approximately 0.5-0.6 kcal/mol. Although the G:s^2^C pair is thermodynamically less stable than A:s^2^U and I:s^2^C, the smaller Δ*G*° penalty associated with dG incorporation results in the dG:s^2^C pair exhibiting comparable stability to dA:s^2^U and dI:s^2^C. As a result, dA:s^2^U, dI:s^2^C, and dG:s^2^C can all be considered isoenergetic at 25°C.

Although the dG:s^2^C and dI:s^2^C base pairs are isoenergetic at 25°C, their stabilities diverge at elevated temperatures due to opposing entropic contributions: dG:s^2^C incurs an entropic penalty, whereas dI:s^2^C benefits from an entropic gain. As a result, dG:s^2^C becomes less stable than dI:s^2^C at higher temperatures. For instance, the Δ*G*° values of the two base pairs differ by approximately 0.6 kcal/mol at 65 °C. This reduced thermal stability suggests that dG:s^2^C containing sequences are more readily denatured at elevated temperatures, which could be advantageous in nonenzymatic replication by facilitating strand separation after completion of primer extension. If this thermodynamic difference accumulates with increasing proportions of dG:s^2^C relative to dI:s^2^C, it could impose a strong selective pressure favoring the transition from dI to dG in early replication systems. Therefore, if deoxyribonucleotides constituted the majority of purine components rather than ribonucleotides, this entropic behavior could help explain the evolutionary replacement of inosine by guanosine in primordial genetic polymers.

To further investigate the thermodynamic implications of replacing G with dG, we examined the wobble base pair G:U and its deoxyribo-purine counterpart, dG:U. The canonical G:U wobble pair shows a Δ*G*°_25°C_ only 0.8 kcal/mol weaker than the standard A:U base pair, providing sufficient stability to participate in RNA folding and secondary structure formation. In contrast, the dG:U pair exhibits a larger Δ*G*°_25°C_ difference, approximately 1.1–1.3 kcal/mol, when compared to the base pairs dA:s^2^U, dI:s^2^C, and dG:s^2^C. This greater instability indicates that, unlike the G:U wobble pair, the dG:U mismatch is unlikely to pose a significant fidelity concern. Furthermore, in a primordial genetic system dominated by 2-thiopyrimidines, the dG:s^2^U mismatch is not expected to be a concern since s^2^U cannot form a wobble pair with G. On the other hand, while previous findings suggest that s^2^U:s^2^U mismatches are not problematic during nonenzymatic RNA replication when competing with ribo-adenosine (38), they could present a fidelity challenge in systems involving deoxyadenosine. The dA:s^2^U pair is only marginally more stable than the s^2^U:s^2^U mismatch, with a Δ*G*°_25°C_ difference of just 0.6 kcal/mol. This small energetic margin implies a limited thermodynamic preference for correct pairing, potentially compromising replication fidelity in prebiotic systems incorporating deoxyribonucleotides. Further experimental studies are required to assess the extent to which this narrow stability difference affects base selection and sequence propagation under prebiotic conditions.

### Crystal Structure Studies of RNA Duplexes Containing Deoxyribo-purines

To better understand why deoxyribo-purine substitutions result in reduced base pair stability, we designed ten 16-mer self-complementary RNA sequences containing 2′-deoxyribo-purines for structural studies. Among these, four sequences (dAU1, dAU2, dAUS1, and dAUS2) contained base pairs involving deoxyadenosine (dA:U and dA:s^2^U), four (dGC1, dGC2, dGCS1, and dGCS2) contained deoxyguanosine base pairs (dG:C and dG:s^2^C), and two (dICS1 and dICS2) included base pairs with deoxyinosine (dI:s^2^C). Previously reported structures of the corresponding ribo-purine sequences provided the basis for comparisons. The sequences used for crystallization are listed in Table S1. All ten oligonucleotides crystallized within 2–3 days at 20 °C under their respective optimal conditions (Table S2), and their structures were solved by X-ray diffraction at resolutions ranging from 1.2 to 2.0 Å. Data collection and refinement statistics are summarized in Tables S3 and S4. Eight of the ten structures crystallized in the same space group (H32) with similar packing, each unit cell containing a single RNA strand. In contrast, sequences containing dG:s^2^C formed distinct crystal lattices. Notably, the dGCS1 structure also crystallized in space group H32, but with a duplex per unit cell instead of a single strand. The dGCS2 duplex crystallized in space group P31, with six strands per unit cell.

No significant differences in hydrogen bonding between nucleobases were observed between ribo-purines and 2′-deoxyribo-purines when forming equivalent base pairs (Figure S3). When paired with the same pyrimidines, including 2-thiopyrimidines, ribo- and deoxyribo-purines formed the same number of hydrogen bonds with similar bond lengths. Additionally, the introduction of a single deoxyribo-purine modification into RNA duplexes did not result in any detectable perturbation of the local sugar pucker conformation or the overall helical geometry.

Due to the absence of the 2′-hydroxyl group, the 2′-deoxyribose sugars exhibit fewer hydrogen bonding interactions with surrounding solvent molecules. Similarly to earlier studies, our structural analyses revealed multiple water molecules forming hydrogen bonds with the ribose units of both purines and pyrimidines (39-41). Notably, one water molecule (Water 1) was consistently observed forming hydrogen bonds with both the 2′- and 3′- hydroxyl groups of most base pairs, including those involving adenosine, guanosine, and inosine (Figures 2A, B, E, F, M, N, S). Water 1 typically forms a shorter hydrogen bond with the 2′-hydroxyl group, with bond lengths ranging from 2.7 to 3.0 Å, and a slightly longer and presumably weaker hydrogen bond with the 3′-hydroxyl group, with bond lengths ranging from 2.9 to 3.5 Å. In contrast, when ribo-purines are replaced by deoxyribo-purines in the same base pairs, the absence of the 2′-hydroxyl group means that Water 1 (Figures 2C, D, G, H, O, T) can only engage with the 3′-hydroxyl group, introducing increased positional flexibility and weakening the interaction, as reflected by a slightly longer hydrogen bond lengths of approximately 2.9 to 3.6 Å. In some cases, likely due to changes in crystal packing, Water 1 is not visible, for example at the dG7 position in dGCS2 (Figure 2P), in contrast to its clear presence at G7 in GCS2 (Figure 2N).

**Figure 1.**
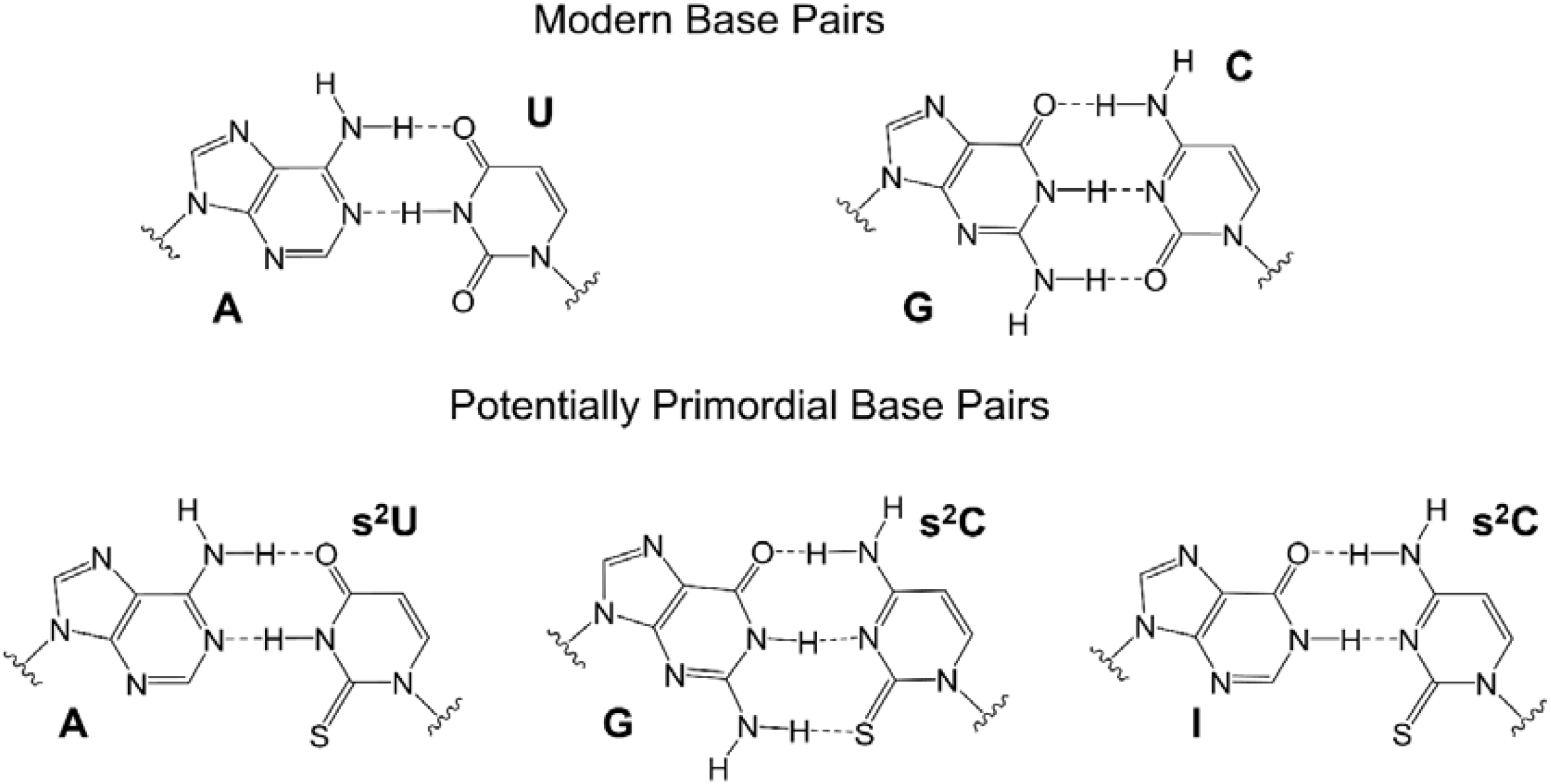
Schematic structures of the modern A:U and G:C base pairs (Top) and potentially primordial base pairs A:s^2^U, G:s^2^C, and I:s^2^C (Bottom).

**Figure 2.**
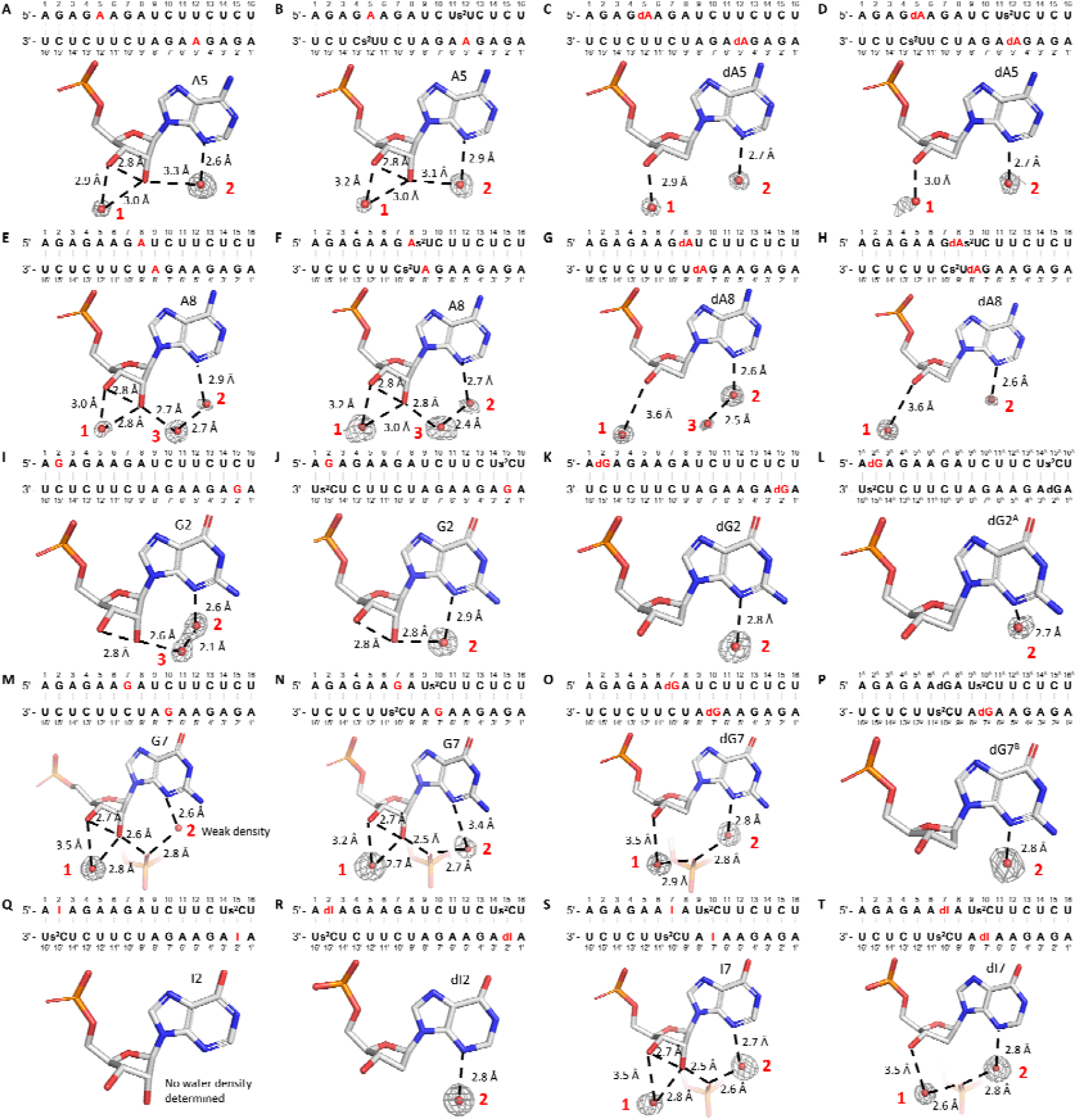
Solvent hydrogen binding interactions of ribo- and deoxyribo-purines in 16mer RNA duplex crystal structures. Sequence context and solvent-mediated hydrogen bonding interactions are shown for the following purine positions: (A) A5 in Native16; (B) A5 in AUS1; (C) dA5 in dAU1; (D) dA5 in dAUS1; (E) A8 in Native16; (F) A8 in AUS2; (G) dA8 in dAU2; (H) dA8 in dAUS2; (I) G2 in Native16; (J) G2 in GCS1; (K) dG2 in dGC1; (L) dG2 in dGCS1; (M) G7 in Native16; (N) G7 in GCS2; (O) dG7 in dGC2; (P) dG7 in dGCS2; (Q) I2 in ICS1; (R) dI2 in dICS1; (S) I7 in ICS2; (T) dI7 in dICS2. Red numbers indicate the number of water molecules. Gray mesh represents the corresponding 2F_o_-F_c_ omit maps for water molecules contoured at 1.0 σ. The structures of Native16, AUS1, AUS2, GCS1, GCS2, ICS1, and ICS2 were previously reported in ref (37), and the figures presented here are derived from the corresponding PDB entries: 9CSO, 9CSP, 9CSQ, 9CSR, 9MDW, 9MDX, and 9MDY, respectively.

Another commonly observed water molecule (Water 2) forms a hydrogen bond with the N3 atom of the purine base. In several ribo-purine containing structures, Water 2 is also hydrogen bonded to the 2′-hydroxyl group (Figures 2A, B, J), with bond lengths ranging from 2.8 to 3.3 Å. This dual interaction enhances local hydration and contributes to the structural stability of the RNA duplex. However, in structures containing 2′-deoxyribo-purine substitutions, although Water 2 maintains its interaction with the N3 atom, loss of the hydrogen bond with the 2′-hydroxyl group (Figures 2C, D, L) may contribute to the overall weaker solvation environment associated with 2′-deoxyribo-purines.

In other cases, Water 2 forms a hydrogen bond with another water molecule (Water 3), which in turn is hydrogen bonded to the 2′-hydroxyl group of the ribo-purine (Figures 2E, F, I). When ribo-purines are replaced by 2′-deoxyribo-purines, Water 3 is typically absent (Figures 2H and K). Even when Water 3 is present, as in Figure 2G, the weak electron density suggests that it is disordered, likely due to the increased flexibility resulting from the loss of hydrogen bonding with the missing 2′-hydroxyl group. In some instances, Water 3 is replaced by a non-bridging oxygen from a phosphate group in a neighboring RNA strand, depending on the crystal packing. This substitution was specifically observed at G7 or I7 positions, reflecting inter-strand interactions in the crystal lattice (Figures 2M, N, S). In 2′-deoxyribo-purine containing base pairs, the non-bridging phosphate oxygen remains present as long as the crystal packing is preserved, but with the loss of one hydrogen bond between the non-bridging oxygen and 2′-hydroxyl group (Figures 2O and T). However, in the case of dG7 in the dGCS2 structure, altered crystal packing prevents Water 2 from forming hydrogen bonds with either Water 3 or non-bridging phosphate oxygens.

The reduced solvation observed for 2′-deoxyribo-purines, along with the loss of stabilizing hydrogen bonds between the 2′- and 3′-hydroxyl groups, may increase the conformational flexibility of the sugar moiety. This enhanced flexibility may contribute to the observed decrease in base pair stability. Notably, sequences containing dG:s^2^C pairs crystallized in a distinct lattice from the same sequence with G:s^2^C pairs, which may reflect a markedly increased flexibility in dG sugar pucker conformation compared to dA or dI. This interpretation is consistent with thermodynamic data showing a greater entropic penalty associated with replacing rG with dG.

### Crystal Structure Studies of RNA Duplexes with RNA or DNA Imidazolium-bridged-dinucleotides

Thus far, our thermodynamic and structural studies have primarily focused on the effects of 2′-deoxyribo-purine substitutions within RNA duplexes. However, these analyses alone are insufficient to fully understand the impact of 2′-deoxyribo-purines on nonenzymatic RNA replication. To gain deeper insight into how DNA-like intermediates might influence the chemistry of nonenzymatic primer extension, we employed a crystal soaking approach—a technique that previously enabled us to directly visualize the imidazolium-bridged dinucleotide intermediate during RNA primer extension (42). In this study, we co-crystallized a 14-mer LNA-modified self-complementary RNA sequence (Figure 3A) in the presence of dGMP, which was positioned on the 2-nucleotide 5′-overhangs at each end of the duplex. Crystals were subsequently soaked in droplets containing either the ribonucleotide substrate guanosine-5′-phosphoro-2-aminoimidazolide (*prG, Figure 3B) or the corresponding substrate 2′-deoxyguanosine-5′-phosphoro-2-aminoimidazolide (*pdG, Figure 3D) to initiate primer extension. Soaking was carried out for 2 hours with *prG and 4 hours with *pdG to allow sufficient time for reaction progression. The crystals were then harvested by flash-freezing in liquid nitrogen and subjected to X-ray diffraction analysis to capture snapshots of the formation of the imidazolium-bridged dinucleotide intermediates (Figures 3C and E).

**Figure 3.**
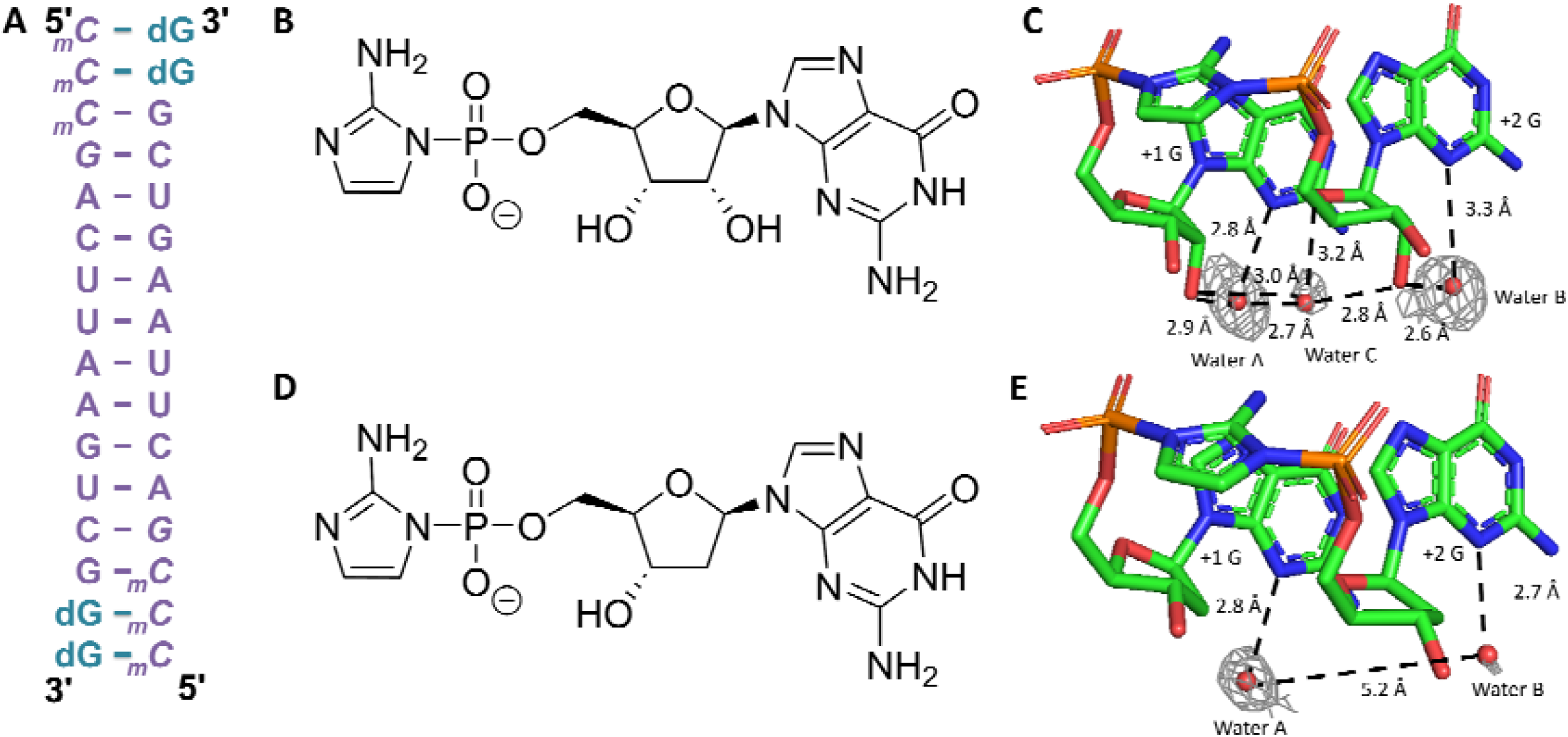
Structure of the imidazolium-bridged rG and dG dinucleotide intermediate bound to an RNA primer/template complex. (A) Schematic of the RNA-dGMP complex used in crystallographic soaking experiments. The four italic nucleotides at the 5′-end represent locked nucleic acid. (B) Chemical structure of guanosine-5′-phosphoro-2-aminoimidazolide (prG). (C) Crystal structure of the imidazolium-bridged riboguanosine dinucleotide intermediate (G*G) bound to the RNA duplex. (D) Chemical structure of deoxyguanosine-5′- phosphoro-2-aminoimidazolide (pdG). (E) Crystal structure of the imidazolium-bridged deoxyguanosine dinucleotide intermediate (dG*dG) bound to the RNA duplex. Gray mesh represents the corresponding 2F_o_–F_c_ omit maps for water molecules contoured at 1.0 σ.

The sugar pucker at both the +1 and +2 positions in the G*G and dG*dG imidazolium-bridged intermediates adopted the 3′-endo conformation, which is consistent with efficient primer extension. In the G*G structure, a network of high occupancy, well-ordered water molecules was identified, contributing to the stabilization of the intermediate. One water molecule (water A) simultaneously coordinates to the N3 position of the +1 G and the 2′- hydroxyl of the same nucleotide. A second water molecule (water B) exhibits a similar pattern, coordinating to the N3 of the +2 G and its corresponding 2′-hydroxyl group. A third water molecule (water C) bridges waters A and B, while also forming a hydrogen bond with the 2′- hydroxyl group of the +1 G and O4′ atom of the +2 G, establishing an extensive and cooperative hydration network that reinforces the structural integrity of the G*G intermediate. In contrast, the dG*dG structure exhibited fewer and less well-ordered solvent-mediated interactions. While water A remains well-defined and forms a similar coordination with N3 of the +1 dG, the density for water B is considerably weaker, likely due to the absence of the 2′-hydroxyl group in deoxyribonucleotides. Moreover, no electron density corresponding to water C was observed, and the distance between water A and B extends to approximately 5.2 Å, suggesting a lack of direct interaction between the two. This reduced hydration network in the dG*dG intermediate may lead to weaker template binding or diminished structural stability relative to its ribonucleotide counterpart.

### Kinetic Study of Primer Extension with Deoxyribo-purines

Because our thermodynamic and structural analyses suggested that 2′-deoxyribo-purines form less stable base pairs than ribo-purines, we proceeded to investigate the kinetic performance of deoxyribo-purines in nonenzymatic primer extension reactions. For these experiments we employed a primer/helper/template system consisting of a 26-nucleotide template hybridized to a 12-nucleotide primer and a 12-nucleotide downstream helper, forming a 2-nucleotide gap in which imidazolium-bridged dinucleotide substrates could bind (Figure 4A and Figure S4). Using this system, we conducted nonenzymatic RNA template copying reactions with complementary substrate-template pairs across a range of substrate concentrations. The resulting kinetic data were analyzed by fitting to the Michaelis–Menten equation (Figure S5), allowing us to extract Michaelis–Menten constants (*K*_m_) and maximal observed reaction rates (*k*_obs max_). These kinetic parameters are summarized and visualized as heatmaps in Figures 4B and C.

**Figure 4.**
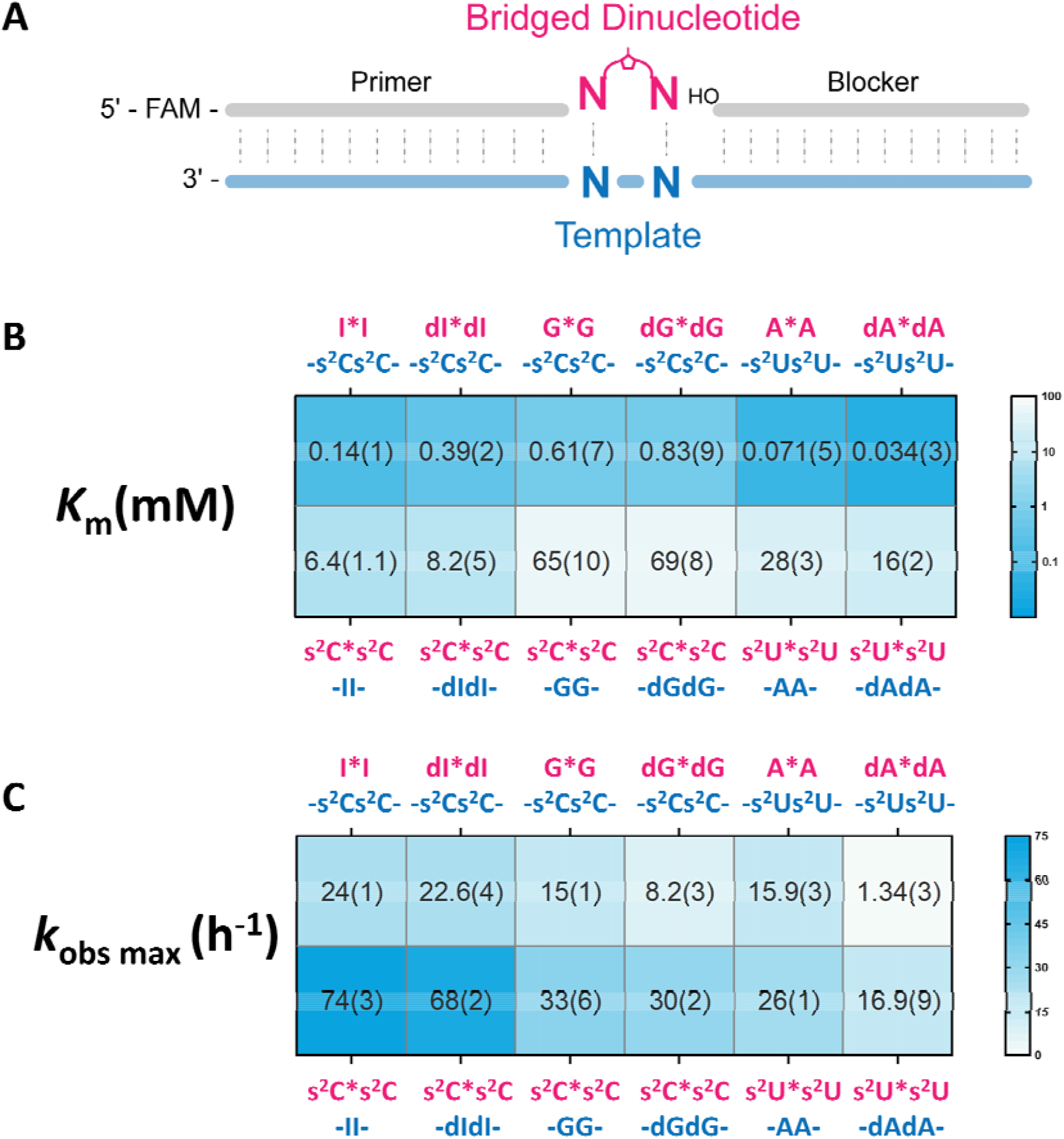
(A) Schematic representation of the nonenzymatic primer extension reactions. (B) Michaelis-Menten constant (K_m_) of the bridged dinucleotides on the template. (C) Observed maximum rate (k_obs max_) of the reaction. All reactions were performed at room temperature with 1.5 µM primer, 2.5 µM template, 3.5 µM blocker, 100 mM MgCl_2_, and 200 mM Tris-HCl pH 8.0. Standard errors (N ≥ 2) are reported in parentheses. The values of s ^2^C*s ^2^C on -dIdI-, dI*dI on -s ^2^Cs ^2^C-, s^2^ C*s^2^ C on -dGdG-, dG*dG on - s^2^ Cs ^2^C -, s ^2^U*s ^2^U on -dAdA-, and dA*dA on -s^2^U*s^2^U- are reported here for the first time. The values for s^2^C*s^2^C on -II-, I*I on -s^2^Cs^2^C-, s^2^C*s^2^C on -GG-, G*G on - s^2^Cs^2^C -, s^2^U*s^2^U on -AA-, and A*A on -s^2^U*s^2^U- are reproduced from Figure 3 in ref (37) with permission under a Creative Commons Attribution 4.0 International License. Copyright 2025 National Academy of Sciences.

The dI*dI imidazolium bridged dinucleotide substrates exhibited moderately reduced binding affinity, compared to its ribo-counterpart (I*I) on an -s^2^Cs^2^C-template, reflected by a 2.8-fold increase in *K*_m_, while maintaining a comparable maximum reaction rate (22.6 h^−1^ for dI*dI versus 24 h^−1^ for I*I). When dI was incorporated into the template instead of the substrate, the differences in substrate binding and reaction rate became even smaller. The - dIdI-template showed nearly identical reaction kinetics and only a modest 1.3-fold increase in *K*_m_ for s^2^C*s^2^C relative to the -II-template. Consequently, deoxyinosine substitution resulted in only a 2-fold reduction in the disparity in binding affinities between purine and pyrimidine substrates (from 46-fold to 21-fold).

The effect of substituting guanosine with deoxyguanosine is even smaller. When using dG*dG imidazolium bridged dinucleotides as substrates for primer extension on an -s^2^Cs^2^C- template, the *K*_m_ is approximately 1.4-fold higher than that of the G*G substrates, accompanied by a maximum reaction rate that is reduced by ∼1.8-fold. When comparing the kinetic parameters for primer extension using s^2^C*s^2^C substrates on templates containing either -dGdG- or -GG-, the differences are negligible for both maximum reaction rate and binding affinity. This less pronounced effect of 2′-deoxyguanosine compared to 2′- deoxyinosine is consistent with our thermodynamic data, which show that substitution of guanosine with 2′-deoxyguanosine results in smaller changes in duplex hybridization free energy than substitution of inosine with 2′-deoxyinosine. As a result, compared to their ribo-counterparts, dI and dG exhibit much closer binding affinities to s^2^C templates in nonenzymatic primer extension, whether acting as substrates or as template bases. However, the disparity in binding affinities between purine and pyrimidine substrates remains largely unresolved, at approximately 83-fold, comparable to the 107-fold disparity observed with ribo-guanosine.

Surprisingly, the replacement of A with dA had quite different effects than observed with G or I. The maximum reaction rate of the dA*dA imidazolium-bridged dinucleotide was reduced by 12-fold relative to A*A, while its affinity for the -s^2^Us^2^U-template was unexpectedly 2.1-fold greater. A similar trend was observed for the s^2^U*s^2^U substrate, which had a 1.5-fold slower reaction rate and a 1.8-fold increase in binding affinity to the -dAdA-template compared to the -AA- template. These results differ from expectations based on thermodynamic base pairing data, but are consistent with previous kinetic observations (10). We suggest that subtle differences in base pairing thermodynamics and geometry between internal sites within a duplex, and the binding of bridged dinucleotide substrates, may significantly affect the kinetics of primer extension. Despite the enhanced affinities resulting from replacing A with dA, the pyrimidine/purine disparity in reaction rates increased from a 390-fold difference between A*A on a -s^2^Us^2^U- template and s^2^U*s^2^U on a -AA- template, to a 470-fold difference.

Overall, substituting purine ribonucleotides with their 2′-deoxyribo counterparts led to mixed effects on template binding. For dA substrates, the affinity for pyrimidine templates increased, whereas dI and dG substrates exhibited modest decreases in affinity. Similarly, when pyrimidine substrates were paired with deoxyribo-purine templates, binding affinities remained largely unchanged for dI and dG, while small increases were observed with dA templates. As a result, substituting purine ribonucleotides with 2′-deoxyribo-purines did not substantially alleviate the large difference in purine vs. pyrimidine substrate affinities during nonenzymatic primer extension.

## DISCUSSION

The “RNA/DNA world” hypothesis has gained attention in recent years, due in part to experimental demonstrations of plausible prebiotic routes to purine 2′-deoxyribonucleosides (19-21). Rather than assuming that DNA emerged as a wholly distinct genetic system from an RNA-dominated world, this model envisions a more gradual transition from an ancestral RNA/DNA mixed world. A mixed nucleic acid system, comprised of both ribonucleotides and deoxyribonucleotides, may have represented a transitional genetic system capable of supporting both primitive information storage and catalysis. Our thermodynamic and structural findings are consistent with a possible role of purine 2′-deoxyribonucleotides in a primordial replication system.

At the same time, our data reveals challenges that complicate this picture. For example, our structural studies of 2′-deoxyribo-purine containing duplexes reveal diminished solvent-mediated hydrogen bonding relative to their ribose counterparts. This suggests that the presence of deoxyribo-purines in functional RNAs could weaken critical secondary structures. Nevertheless, the discovery and characterization of functional DNAzymes demonstrates that catalysis is not uniquely dependent on the ribose sugar (43). Moreover, our previous work and that of others have shown that nonheritable backbone heterogeneity can be tolerated in primitive functional systems (44). This flexibility implies that early nucleic acid pools may not have required a uniform sugar chemistry to support sequence evolution, replication, and function. *In vitro* selection of novel chimeric ribozymes from randomized sequence libraries could be used to test the importance of the heritability of sugar chemistry, for example by comparing libraries containing both ribonucleotides and deoxyribonucleotides, in which the sites of sugar heterogeneity are not heritable, and libraries consisting of pyrimidine ribonucleotides and purine 2′-deoxyribonucleotides, in which the sugar differences would be heritable. Such studies could determine whether functionally active ribozymes or ligases can emerge from hybrid systems, offering direct evidence of the compatibility or lack thereof between structural heterogeneity and biological function.

From a structural perspective, our crystallographic analyses reveal that the incorporation of one or two deoxyribo-purine substitutions into RNA duplexes does not cause significant perturbations in either the local sugar pucker or the overall helical geometry. However, the cumulative effects of multiple deoxyribonucleotide substitutions remain inadequately characterized. Prior studies have demonstrated that extensive incorporation of deoxyribonucleotides can shift the sugar pucker equilibrium and alter essential helical parameters (45), which are critical for efficient template-directed synthesis in nonenzymatic replication. Because nonenzymatic RNA primer extension depends on the 3′-endo sugar conformation and A-form helical geometry to ensure optimal substrate alignment and catalytic efficiency, deviations toward DNA-like (C2′-endo) conformations could negatively impact both fidelity and reaction kinetics (46,47). Therefore, additional structural and kinetic investigations of multiply modified RNA/DNA hybrid duplexes are essential for assessing their potential to support prebiotic replication processes.

Another significant concern is the effect of specific mismatches on fidelity. While our earlier studies demonstrated that s^2^U:s^2^U mismatches do not significantly compromise the fidelity of nonenzymatic RNA replication in the presence of sufficient competing adenosine substrates (38), the situation would be less favorable if adenosine was replaced by deoxyadenosine. The dA:s^2^U base pair is only slightly more stable than the s^2^U:s^2^U mismatch, with a Δ*G*° difference of just 0.6 kcal/mol at 25 °C. This small thermodynamic difference suggests that incorrect s^2^U:s^2^U pairings would increase the probability of misincorporation and thus compromise replication fidelity in primordial systems enriched with 2′-deoxyribo-purines. To better quantify this effect, competition experiments involving dA, dI, dG, s^2^U, and s^2^C as substrates copying s^2^U-containing templates would provide insights into how these mismatches impact fidelity.

A final challenge arising from the incorporation of purine deoxyribonucleotides into the primordial genetic alphabet is their effect on the overall efficiency of nonenzymatic primer extension. Our experiments show primer extension efficiency is significantly impaired because dI and dG substrates exhibit reduced binding affinities, while dA substrates suffer from markedly slower reaction kinetics. Notably, their presence did not alleviate the inherent kinetic disparity between purine and pyrimidine substrates. Nevertheless, it is important to consider the broader chemical context of early Earth. The proposed prebiotic synthesis of 2′- deoxyribo-purines is considerably more complex than that of 2-thiopyrimidines (19), suggesting that their concentrations may have been substantially lower in primordial environments. In such scenarios, the kinetic imbalances between purines and pyrimidines observed in our experiments could have been mitigated by differential substrate abundances rather than by intrinsic chemical properties.

Taken together, our results both support and complicate the “RNA/DNA world” hypothesis. They suggest that while deoxyribo-purines may have been viable components of early genetic systems, their integration likely imposed trade-offs between stability, fidelity, and catalytic potential. Continued experiments that model replication, selection, and catalysis in mixed backbone systems will be essential to clarify the viability of this evolutionary pathway.

## DATA AVAILABILITY

Atomic coordinates and structure factors for the reported crystal structures have been deposited with the Protein Data bank under accession numbers 9OKS, 9OKT, 9OKU, 9OKV, 9OKW, 9OKX, 9OKY, 9OKZ, 9OL0, 9OL1, 9OL2, 9OL3.

## SUPPLEMENTARY DATA

Supplementary Data are available at NAR online.

## ACKNOWLEDGEMENTS

The authors thank Dr. Benjamin Colville, Dr. Longfei Wu, and other members of the Szostak laboratory for helpful discussions.

The authors thank the staff at the Advanced Light Source (ALS) beamline 501, 503, 821, National Synchrotron Light Source II (NSLS-II) beamline 17ID-2 and the Advanced Photon Source (APS) beamline 23ID-B.

## REFERENCES

1. Gilbert, W. (1986) Origin of life: The RNA world. Nature, 319, 618–618.

2. Joyce, G.F. (2002) The antiquity of RNA-based evolution. Nature, 418, 214–221.

3. Kozlov, I.A., Politis, P.K., Van Aerschot, A., Busson, R., Herdewijn, P. and Orgel, L.E. (1999) Nonenzymatic synthesis of RNA and DNA oligomers on hexitol nucleic acid templates: the importance of the A structure. Journal of the American Chemical Society, 121, 2653–2656.

4. Gavette, J.V., Stoop, M., Hud, N.V. and Krishnamurthy, R. (2016) RNA–DNA chimeras in the context of an RNA world transition to an RNA/DNA world. Angewandte Chemie International Edition, 55, 13204–13209.

5. Orgel, L. (2000) A simpler nucleic acid. Science, 290, 1306–1307.

6. Chaput, J.C. (2021) Redesigning the genetic polymers of life. Accounts of Chemical Research, 54, 1056–1065.

7. Zhang, W., Kim, S.C., Tam, C.P., Lelyveld, V.S., Bala, S., Chaput, J.C. and Szostak, J.W. (2021) Structural interpretation of the effects of threo-nucleotides on nonenzymatic template-directed polymerization. Nucleic acids research, 49, 646–656.

8. Taylor, A.I., Pinheiro, V.B., Smola, M.J., Morgunov, A.S., Peak-Chew, S., Cozens, C., Weeks, K.M., Herdewijn, P. and Holliger, P. (2015) Catalysts from synthetic genetic polymers. Nature, 518, 427–430.

9. Roberts, S.J., Szabla, R., Todd, Z.R., Stairs, S., Bučar, D.-K., Šponer, J., Sasselov, D.D. and Powner, M.W. (2018) Selective prebiotic conversion of pyrimidine and purine anhydronucleosides into Watson-Crick base-pairing arabino-furanosyl nucleosides in water. Nature Communications, 9, 4073.

10. Kim, S.C., Zhou, L., Zhang, W., O’Flaherty, D.K., Rondo-Brovetto, V. and Szostak, J.W. (2020) A model for the emergence of RNA from a prebiotically plausible mixture of ribonucleotides, arabinonucleotides, and 2′-deoxynucleotides. Journal of the American Chemical Society, 142, 2317–2326.

11. Joyce, G.F. (2012) Toward an alternative biology. Science, 336, 307–308.

12. Eschenmoser, A. (1999) Chemical etiology of nucleic acid structure. Science, 284, 2118–2124.

13. Kim, S.C., O’Flaherty, D.K., Giurgiu, C., Zhou, L. and Szostak, J.W. (2021) The emergence of RNA from the heterogeneous products of prebiotic nucleotide synthesis. Journal of the American Chemical Society, 143, 3267–3279.

14. Oró, J. and Stephen-Sherwood, E. (1974) The prebiotic synthesis of oligonucleotides. Origins of life, 5, 159–172.

15. Benner, S.A., Ellington, A.D. and Tauer, A. (1989) Modern metabolism as a palimpsest of the RNA world. Proceedings of the National Academy of Sciences, 86, 7054–7058.

16. Reichard, P. (1993) From RNA to DNA, why so many ribonucleotide reductases? Science, 260, 1773–1777.

17. Leu, K., Obermayer, B., Rajamani, S., Gerland, U. and Chen, I.A. (2011) The prebiotic evolutionary advantage of transferring genetic information from RNA to DNA. Nucleic acids research, 39, 8135–8147.

18. Bhowmik, S. and Krishnamurthy, R. (2019) The role of sugar-backbone heterogeneity and chimeras in the simultaneous emergence of RNA and DNA. Nature chemistry, 11, 1009–1018.

19. Xu, J., Chmela, V., Green, N.J., Russell, D.A., Janicki, M.J., Góra, R.W., Szabla, R., Bond, A.D. and Sutherland, J.D. (2020) Selective prebiotic formation of RNA pyrimidine and DNA purine nucleosides. Nature, 582, 60–66.

20. Xu, J., Green, N.J., Gibard, C., Krishnamurthy, R. and Sutherland, J.D. (2019) Prebiotic phosphorylation of 2-thiouridine provides either nucleotides or DNA building blocks via photoreduction. Nature Chemistry, 11, 457–462.

21. Xu, J., Green, N.J., Russell, D.A., Liu, Z. and Sutherland, J.D. (2021) Prebiotic photochemical coproduction of purine ribo-and deoxyribonucleosides. Journal of the American Chemical Society, 143, 14482–14486.

22. Nordlund, P. and Reichard, P. (2006) Ribonucleotide reductases. Annu. Rev. Biochem., 75, 681–706.

23. Lundin, D., Berggren, G., Logan, D.T. and Sjöberg, B.-M. (2015) The origin and evolution of ribonucleotide reduction. Life, 5, 604–636.

24. Chheda, U., Pradeepan, S., Esposito, E., Strezsak, S., Fernandez-Delgado, O. and Kranz, J. (2024) Factors affecting stability of RNA–temperature, length, concentration, pH, and buffering species. Journal of Pharmaceutical Sciences, 113, 377–385.

25. Fang, Z., Pazienza, L.T., Zhang, J., Tam, C.P. and Szostak, J.W. (2024) Catalytic Metal Ion–Substrate Coordination during Nonenzymatic RNA Primer Extension. Journal of the American Chemical Society, 146, 10632–10639.

26. Bashkin, J.K. and Jenkins, L.A. (1994) The role of metals in the hydrolytic cleavage of DNA and RNA. Comments on Inorganic Chemistry, 16, 77–93.

27. Zhou, L., Ding, D. and Szostak, J.W. (2021) The virtual circular genome model for primordial RNA replication. Rna, 27, 1–11.

28. Ding, D., Zhou, L., Mittal, S. and Szostak, J.W. (2023) Experimental tests of the virtual circular genome model for nonenzymatic RNA replication. Journal of the American Chemical Society, 145, 7504–7515.

29. Ding, D., Zhou, L., Giurgiu, C. and Szostak, J.W. (2022) Kinetic explanations for the sequence biases observed in the nonenzymatic copying of RNA templates. Nucleic Acids Research, 50, 35–45.

30. Otwinowski, Z. and Minor, W. (1997), Methods in enzymology. Elsevier, Vol. 276, pp. 307–326.

31. Kabsch, W. (2010) XDS. Acta Crystallographica Section D: Biological Crystallography, 66, 125–132.

32. McCoy, A.J., Grosse-Kunstleve, R.W., Adams, P.D., Winn, M.D., Storoni, L.C. and Read, R.J. (2007) Phaser crystallographic software. Journal of applied crystallography, 40, 658–674.

33. Mooers, B.H. and Singh, A. (2011) The crystal structure of an oligo (U): pre-mRNA duplex from a trypanosome RNA editing substrate. Rna, 17, 1870–1883.

34. Liebschner, D., Afonine, P.V., Baker, M.L., Bunkóczi, G., Chen, V.B., Croll, T.I., Hintze, B., Hung, L.-W., Jain, S. and McCoy, A.J. (2019) Macromolecular structure determination using X-rays, neutrons and electrons: recent developments in Phenix. Acta Crystallographica Section D: Structural Biology, 75, 861–877.

35. Murshudov, G.N., Vagin, A.A. and Dodson, E.J. (1997) Refinement of macromolecular structures by the maximum-likelihood method. Acta Crystallographica Section D: Biological Crystallography, 53, 240–255.

36. Emsley, P., Lohkamp, B., Scott, W.G. and Cowtan, K. (2010) Features and development of Coot. Acta Crystallographica Section D: Biological Crystallography, 66, 486–501.

37. Fang, Z., Jia, X., Xing, Y. and Szostak, J.W. (2025) Nonenzymatic RNA copying with a potentially primordial genetic alphabet. Proceedings of the National Academy of Sciences, 122, e2505720122.

38. Ding, D., Fang, Z., Kim, S.C., O’Flaherty, D.K., Jia, X., Stone, T.B., Zhou, L. and Szostak, J.W. (2024) Unusual base pair between two 2-thiouridines and its implication for nonenzymatic RNA copying. Journal of the American Chemical Society, 146, 3861–3871.

39. Auffinger, P. and Westhof, E. (1997) Rules governing the orientation of the 2′-hydroxyl group in RNA. Journal of molecular biology, 274, 54–63.

40. Sundaralingam, M. and Pan, B. (2002) Hydrogen and hydration of DNA and RNA oligonucleotides. Biophysical chemistry, 95, 273–282.

41. Fohrer, J., Hennig, M. and Carlomagno, T. (2006) Influence of the 2′-hydroxyl group conformation on the stability of A-form helices in RNA. Journal of Molecular Biology, 356, 280–287.

42. Zhang, W., Walton, T., Li, L. and Szostak, J.W. (2018) Crystallographic observation of nonenzymatic RNA primer extension. Elife, 7, e36422.

43. Santoro, S.W. and Joyce, G.F. (1997) A general purpose RNA-cleaving DNA enzyme. Proceedings of the national academy of sciences, 94, 4262–4266.

44. Trevino, S.G., Zhang, N., Elenko, M.P., Lupták, A. and Szostak, J.W. (2011) Evolution of functional nucleic acids in the presence of nonheritable backbone heterogeneity. Proceedings of the national academy of sciences, 108, 13492–13497.

45. Shaw, N.N. and Arya, D.P. (2008) Recognition of the unique structure of DNA: RNA hybrids. Biochimie, 90, 1026–1039.

46. Giurgiu, C., Fang, Z., Aitken, H.R., Kim, S.C., Pazienza, L., Mittal, S. and Szostak, J.W. (2021) Structure–Activity Relationships in Nonenzymatic Template□Directed RNA Synthesis. Angewandte Chemie International Edition, 60, 22925–22932.

47. Park, S.J., Callaghan, K.L. and Ellis, A.V. (2023) Role of helicity in the nonenzymatic template-directed primer extension of DNA. Organic & Biomolecular Chemistry, 21, 6702–6706.

