## Supplementary Data for "Impact of 2′-deoxyribo-purine substrates on nonenzymatic RNA template-directed primer extension"

Present Address:

1. **Synthesis and characterization of homo-bridged-dinucleotides**

The 5′-5′-2-aminoimidazolium-bridged-dinucleotides were synthesized following a previously reported procedure (1,2). Among them, A*A, C*C, G*G, U*U, I*I, s^2^C*s^2^C, s^2^U*s^2^U have been described in prior publications (2), whereas dI*dI, dA*dA and dG*dG were synthesized for the first time in this study. Details of the nuclear magnetic resonance (NMR) spectra and high-resolution mass spectrometry (HRMS) of dI*dI, dA*dA and dG*dG are provided below.

**1.1 1,3-di-(3’-deoxyinosine-5ʹphosphoryl)-2-aminoimidazolium (dI*dI)**

**^1^H NMR** (600 MHz, D_2_O) δ 8.19 (s, 2H), 8.15 (s, 2H), 6.65 (s, 2H), 6.34 (t, *J* = 6.5 Hz, 2H), 4.64 (m, 2H), 4.13-4.05 (m, 4H), 4.05-4.00 (m, 2H), 2.79 (m, 2H), 2.54 (m, 2H). Peaks corresponding to residual TEAB observed at 3.21 and 1.29 ppm.

**^31^P NMR** (162 MHz, D_2_O) δ -12.91. (The purity is 100% with no detectable activated monomer peak)

**HRMS** (Q-TOF) m/z: [M − H]^–^ Calcd. for C_23_H_26_N_11_O_12_P_2_ 710.1238; Found: 710.1307.

**1.2 1,3-di-(3’-deoxyadenosine-5ʹphosphoryl)-2-aminoimidazolium (dA*dA)**

**^1^H NMR** (600 MHz, D_2_O) δ 8.12 (s, 2H), 8.10 (s, 2H), 6.65 (s, 2H), 6.29 (t, *J* = 6.5 Hz, 2H), 4.59 (m, 2H), 4.12-4.06 (m, 4H), 4.01-3.96 (m, 2H), 2.70 (m, 2H), 2.53 (m, 2H). Peaks corresponding to residual TEAB observed at 3.21 and 1.29 ppm.

**^31^P NMR** (162 MHz, D_2_O) δ -12.80. (The purity is 100% with no detectable activated monomer peak)

**HRMS** (Q-TOF) m/z: [M − H]^–^ Calcd. for C_23_H_28_N_13_O_10_P_2_ 708.1558; Found: 708.1685.

**1.3 1,3-di-(3’-deoxyguanosine-5ʹphosphoryl)-2-aminoimidazolium (dG*dG)**

**^1^H NMR** (600 MHz, D_2_O) δ 7.86 (s, 2H), 6.62 (s, 2H),  6.16 (t, *J* = 6.7 Hz, 2H), 4.61 (m, 2H), 4.1-4.05 (m, 4H), 3.98 (m, 2H), 2.74 (m, 2H), 2.47 (m, 2H). Peaks corresponding to residual TEAB observed at 3.21 and 1.29 ppm.

**^31^P NMR** (162 MHz, D_2_O) δ -12.80. (The purity is 100% with no detectable activated monomer peak)

**HRMS** (Q-TOF) m/z: [M − H]^–^ Calcd. for C_23_H_28_N_13_O_12_P_2_ 740.1456; Found: 740.1551.

2. **Supplementary Figures and Tables**

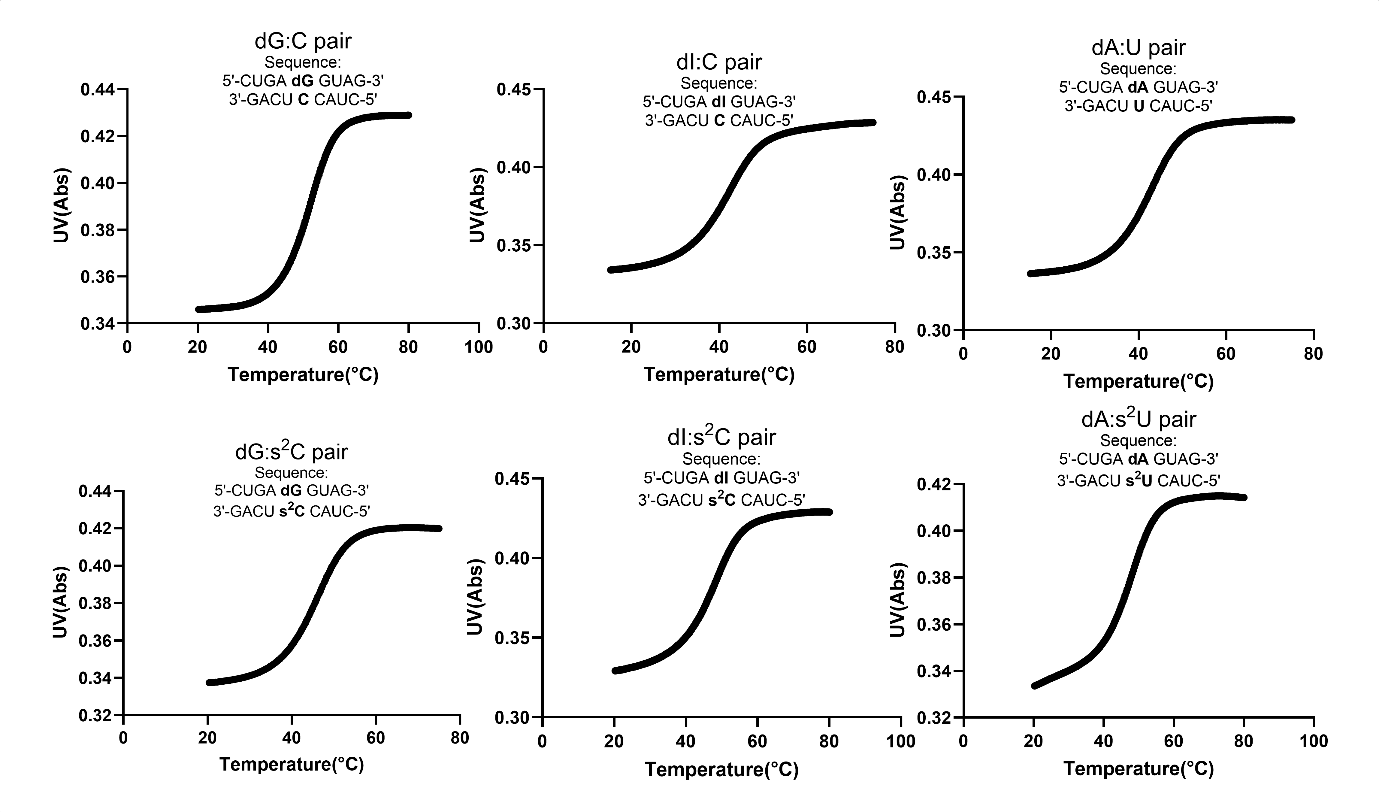

**Figure S1.** Representative melting curves collected during the thermal denaturation of 5 µM oligonucleotide in 10 mM Tris-HCl 8.0, 1 M NaCl, and 2.5 mM EDTA.

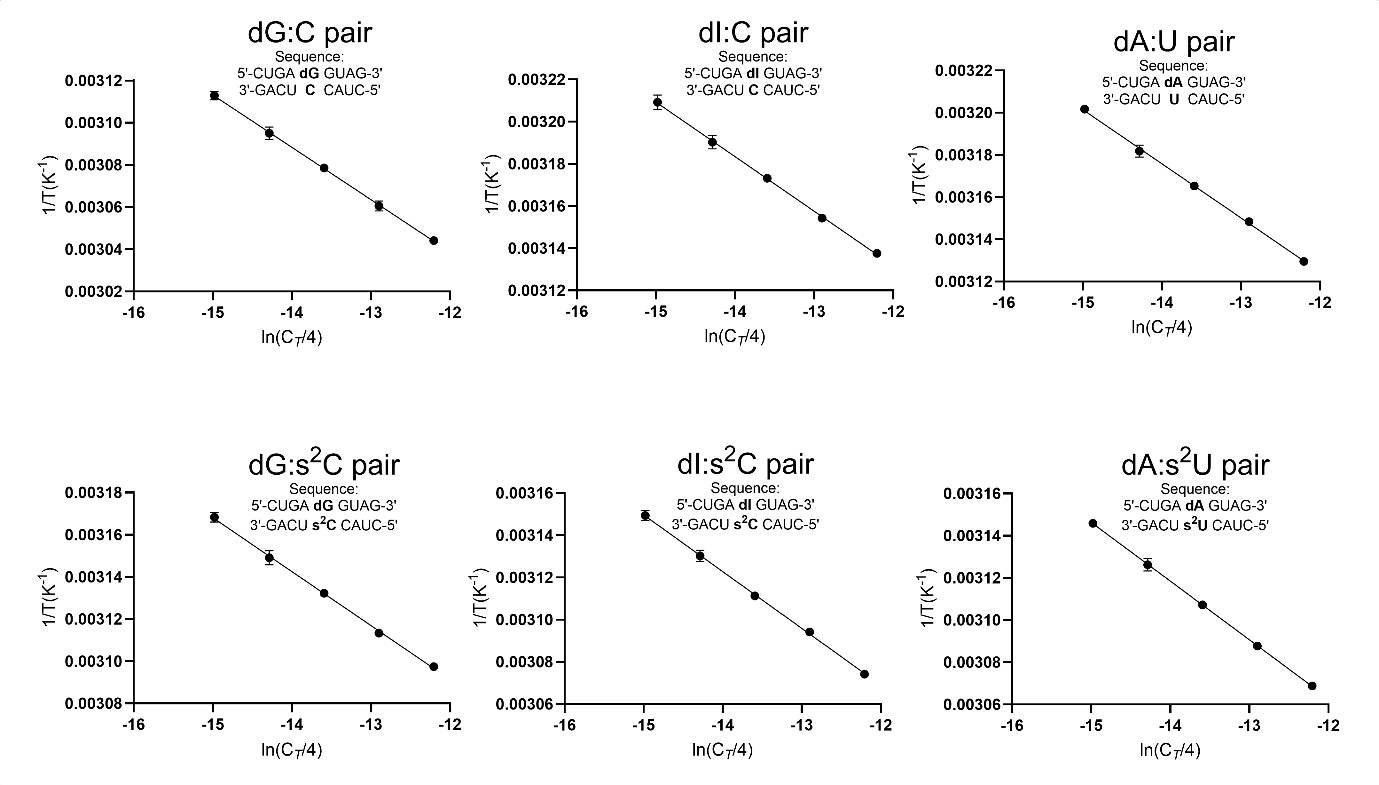

**Figure S2.** Linear least-squared fits of a Van’t Hoff plot of inverse melting temperature (*T*_m_^-1^) collected from optical melts at different oligonucleotide concentrations.

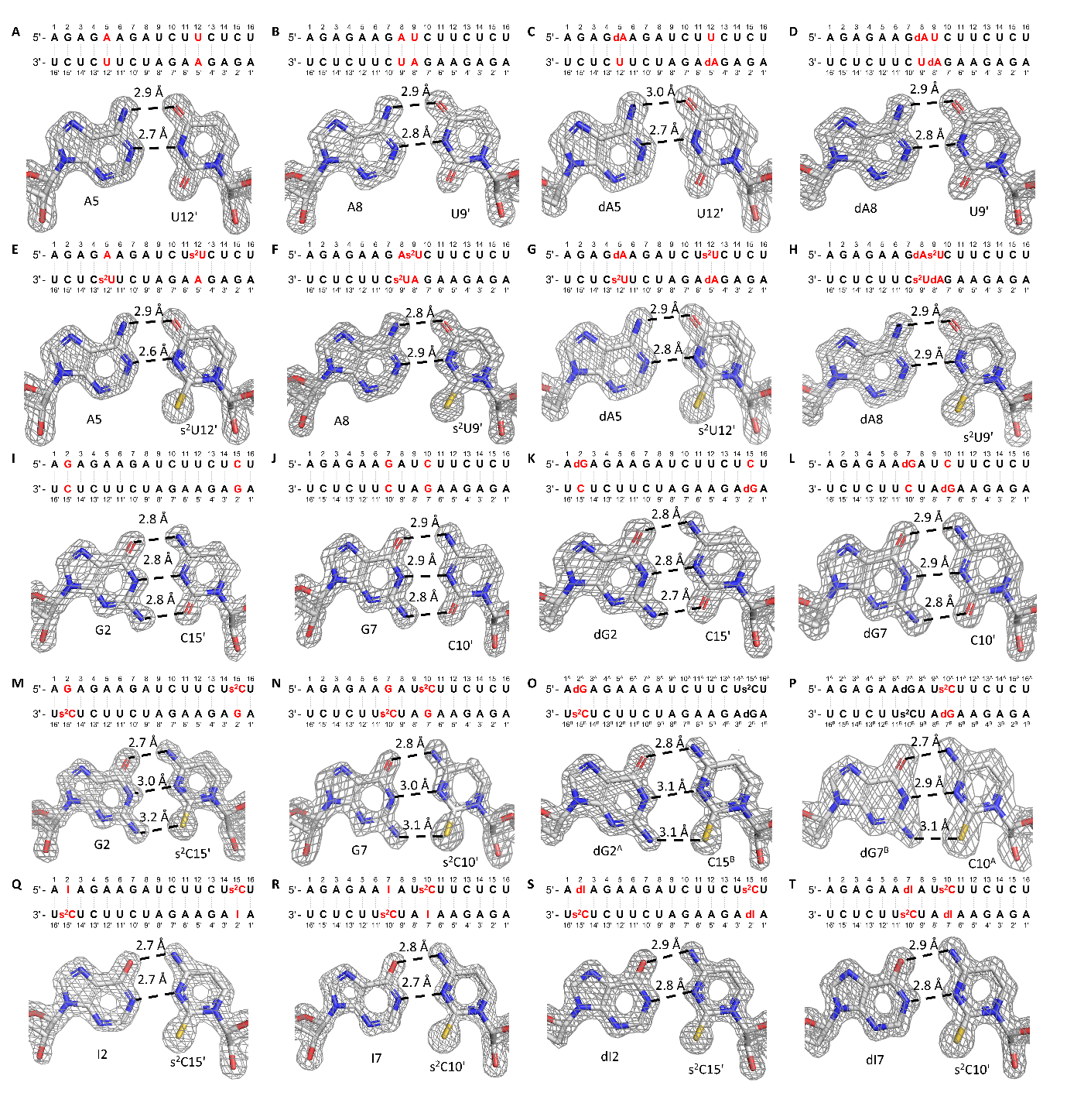

**Figure S3.** Ribo- and deoxy-purines involved base pairs in 16mer RNA duplex crystal structures. Sequence context and density maps are shown for the following base pairs: (A) A5:U12 in Native16; (B) A8:U9 in Native16; (C) dA5:U12 in dAU1; (D) dA8:U9 in in dAU2; (E) A5:s^2^U12 in AUS1; (F) A8:s^2^U9 in AUS2; (G) dA5:s^2^U12 in dAUS1; (H) dA8:s^2^U9 in dAUS2; (I) G2:C15 in Native16; (J) G7:C10 in Native16; (K) dG2:C15 in dGC1; (L) dG7:C10 in dGC2; (M) G2:s^2^C15 in GCS1; (N) G7:s^2^C10 in GCS2; (O) dG2^A^:s^2^C15^B^ in dGCS1; (P) dG7^B^:s^2^C10^A^ in dGCS2; (Q) I2:s^2^C15 in ICS1; (R) I7:s^2^C10 in ICS2; (S) dI2:s^2^C15 in dICS1; (T) dI7:s^2^C10 in dICS2. Gray mesh represents the corresponding 2F_o_-F_c_ omit maps for water molecules contoured at 1.5 σ. The structures of Native16, AUS1, AUS2, GCS1, GCS2, ICS1, and ICS2 were previously reported in reference (2), and the figures presented here are derived from the corresponding PDB entries: 9CSO, 9CSP, 9CSQ, 9CSR, 9MDW, 9MDX, and 9MDY, respectively.

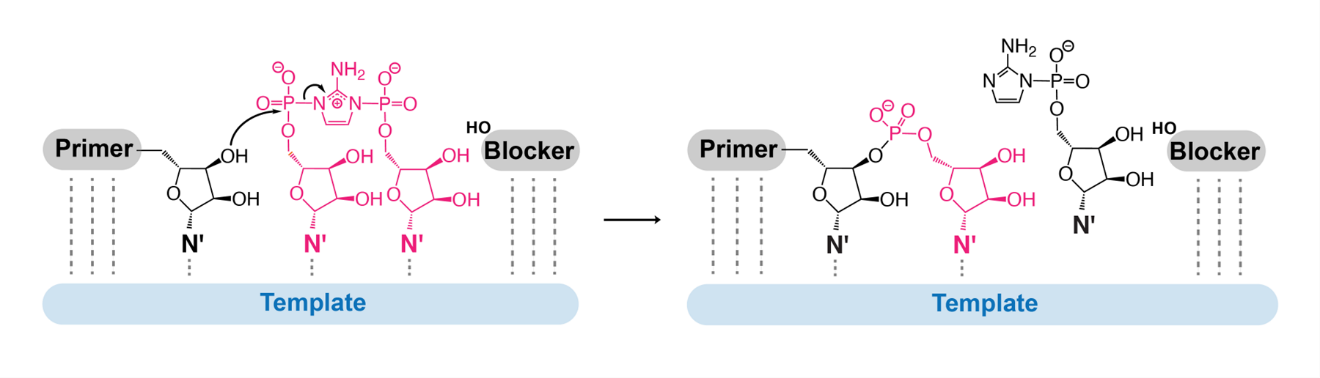

**Figure S4.** Mechanism of bridged dinucleotide (N*N) primer extension within the template-primer-blocker complex

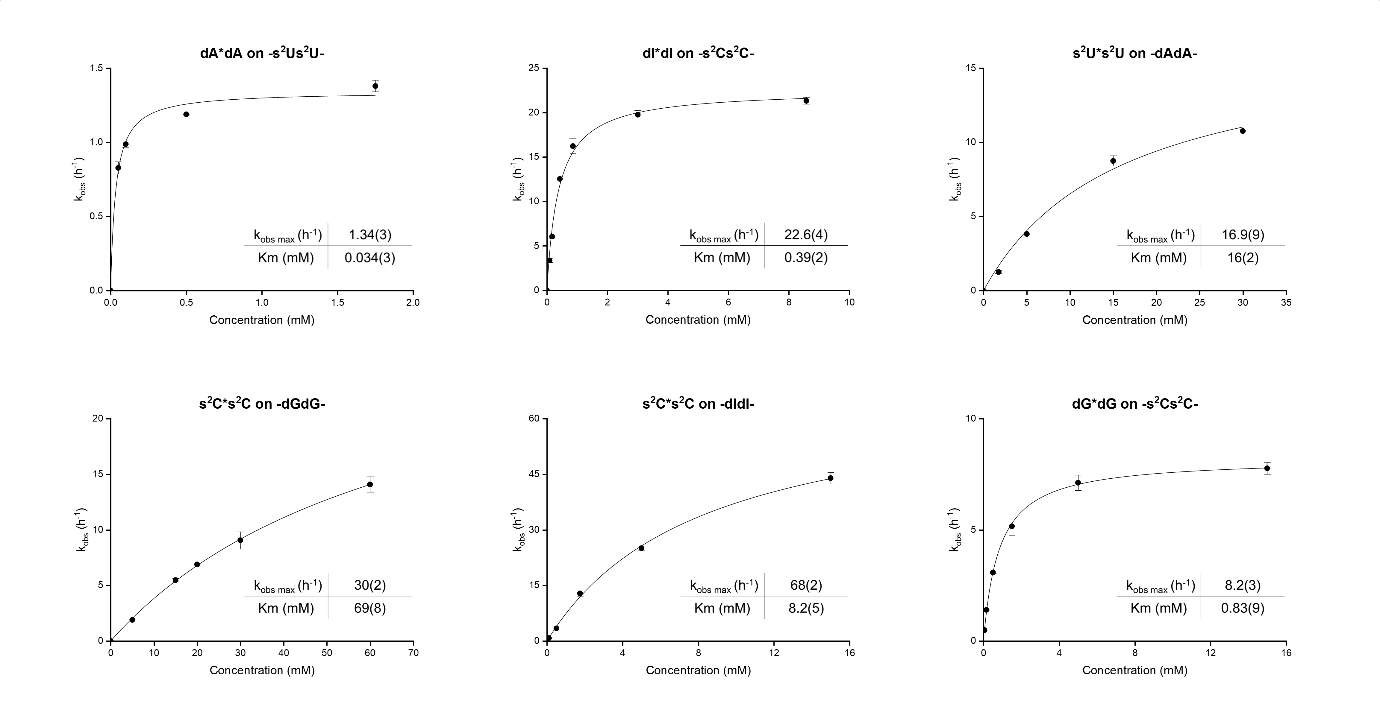

**Figure S5.** Michaelis-Menten curves for primer extension reactions with bridged substrates on the indicated template sequences.

**Table S1.** Oligo Sequences for Crystallization

| Oligo | Sequence |
| --- | --- |
| Native16 | 5’-AGA GAA GAU CUU CUC U-3’ |
| dAU1 | 5’-AGA G**dA**A GAU CU**U** CUC U-3’ |
| dAU2 | 5’-AGA GAA G**dAU** CUU CUC U-3’ |
| AUS1 | 5’-AGA G**A**A GAU CU**s^2^U** CUC U-3’ |
| AUS2 | 5’-AGA GAA G**As^2^U** CUU CUC U-3’ |
| dAUS1 | 5’-AGA G**dA**A GAU CU**s^2^U** CUC U-3’ |
| dAUS2 | 5’-AGA GAA G**dAs^2^U** CUU CUC U-3’ |
| dGC1 | 5’-A**dG**A GAA GAU CUU CU**C** U-3’ |
| dGC2 | 5’-AGA GAA **dG**AU **C**UU CUC U-3’ |
| GCS1 | 5’-A**G**A GAA GAU CUU CU**s^2^C** U-3′ |
| GCS2 | 5’-AGA GAA **G**AU **s^2^C**UU CUC U-3’ |
| dGCS1 | 5’-A**dG**A GAA GAU CUU CU**s^2^C** U-3’ |
| dGCS2 | 5’-AGA GAA **dG**AU **s^2^C**UU CUC U-3’ |
| ICS1 | 5’-A**I**A GAA GAU CUU CU**s^2^C** U-3’ |
| ICS2 | 5’-AGA GAA **I**AU **s^2^C**UU CUC U-3’ |
| dICS1 | 5’-A**dI**A GAA GAU CUU CU**s^2^C** U-3’ |
| dICS2 | 5’-AGA GAA **dI**AU **s^2^C**UU CUC U-3’ |
| 14mer | 5’-*_m_C_m_C_m_C G*AC UUA AGU CG-3’ |

**Table S2.** Optimized Conditions for Crystallization

| Oligo | Optimized crystallization conditions |
| --- | --- |
| dAU1 | 0.08 M Sodium chloride, 0.012 M Potassium chloride, 0.02 M Magnesium chloride hexahydrate, 0.04 M Sodium cacodylate trihydrate pH 6.0, 30% v/v (+/-)-2-Methyl-2,4-pentanediol, 0.012 M Spermine tetrahydrochloride. |
| dAU2 | 0.08 M Sodium chloride, 0.02 M Magnesium chloride hexahydrate, 0.04 M Sodium cacodylate trihydrate pH 5.5, 35% v/v (+/-)-2-Methyl-2,4-pentanediol, 0.002 M Hexammine cobalt(III) chloride. |
| dAUS1 | 0.2 M Magnesium chloride hexahydrate, 0.1 M HEPES sodium pH 7.5, 30% v/v Polyethylene glycol 400. |
| dAUS2 | 0.02 M Calcium chloride dihydrate, 0.1 M Sodium acetate trihydrate pH 4.6, 30% v/v (+/-)-2-Methyl-2,4-pentanediol. |
| dGC1 | 25 % v/v Polyethylene glycol monomethyl ether 550, 50 mM HEPES pH 7.0, 10 mM Magnesium chloride |
| dGC2 | 0.4 M Sodium chloride, 0.12 M Calcium chloride, 20 mM MES pH 5.8, 27% v/v (+/-)-2-Methyl-2,4-pentanediol. |
| dGCS1 | 0.1 M BIS-TRIS pH 5.5, 2.0 M Ammonium sulfate. |
| dGCS2 | 0.1 M MES pH 6.5, 0.06 M Manganese (II) chloride, 15.00 % w/v Polyethylene glycol 20,000 |
| dICS1 | 0.2 M Magnesium chloride hexahydrate, 0.1 M HEPES sodium pH 7.5, 30% v/v Polyethylene glycol 400. |
| dICS2 | 0.2 M Calcium chloride dihydrate, 0.05 M HEPES sodium pH 7.5, 28% v/v Polyethylene glycol 400, 0.002 M Spermine |
| RNAsub | 0.05 M HEPES pH 7.0, 0.2 M Ammonium acetate, 0.15 M Magnesium acetate, 10% w/v Polyethylene glycol 6,000. |
| DNAsub | 0.05 M HEPES pH 7.0, 0.2 M Ammonium acetate, 0.15 M Magnesium acetate, 10% w/v Polyethylene glycol 6,000. |

**Table S3.** Data Collection Statistics

| Sequences | dAU1 | dAU2 | | dAUS1 | dAUS2 |
| --- | --- | --- | --- | --- | --- |
| PDB code | 9OKS | 9OKT | | 9OKU | 9OKV |
| Beamline | ALS 5.0.1 | ALS 5.0.1 | | ALS 5.0.1 | ALS 5.0.1 |
| Wavelength (Å) | 0.97741 | 0.97741 | | 0.97741 | 0.97741 |
| Space group | H32 | H32 | | H32 | H32 |
| Unit cell parameters (Å, °) | 41.2, 41.2, 125.6,  90, 90, 120 | 41.0, 41.0, 123.4,  90, 90, 120 | | 41.2, 41.2, 124.6,  90, 90, 120 | 41.1, 41.1, 124.3,  90, 90, 120 |
| Resolution range (Å) | 50.0-1.50 (1.53-1.50) | 50.0-1.32 (1.34-1.32) | | 50.0-1.48 (1.51-1.48) | 50.0-1.23 (1.25-1.23) |
| Unique reflections | 6909 (334) | 9461 (417) | | 7122 (344) | 11714 (388) |
| Completeness (%) | 100 (100) | 96.9 (88.7) | | 100 (100) | 96.2 (62.3) |
| R_merge_ (%) | 6.7 (51.0) | 6.0 (51.1) | | 6.1 (57.3) | 4.0 (21.4) |
| <I/σ(I)> | 30.0 (4.0) | 30.3 (4.0) | | 33.0 (3.5) | 52.0 (5.2) |
| Redundancy | 8.7 (8.2) | | 9.4 (8.6) | 9.1 (8.6) | 8.3 (3.4) |

| Sequences | dGC1 | dGC2 | dGCS1 | dGCS2 |
| --- | --- | --- | --- | --- |
| PDB code | 9OKW | 9OKX | 9OKY | 9OKZ |
| Beamline | ALS 5.0.3 | ALS 5.0.3 | ALS 5.0.3 | ALS 8.2.1 |
| Wavelength (Å) | 0.97648 | 0.97648 | 0.97648 | 1.00003 |
| Space group | H32 | H32 | H32 | P31 |
| Unit cell parameters (Å, °) | 41.3, 41.3, 124.6,  90, 90, 120 | 41.5, 41.5, 124.6,  90, 90, 120 | 43.4, 43.4, 256.4,  90, 90, 120 | 42.5, 42.5, 122.7,  90, 90, 120 |
| Resolution range (Å) | 50.0-1.54 (1.57-1.54) | 50.0-1.38 (1.40-1.38) | 50.0-1.48 (1.51-1.48) | 50.0-1.92 (1.95-1.92) |
| Unique reflections | 6327 (294) | 8922 (405) | 15145 (756) | 18877 (919) |
| Completeness (%) | 99.9 (100) | 99.5 (96.2) | 93.2 (93.6) | 99.3 (99.4) |
| R_merge_ (%) | 8.7 (33.5) | 6.7 (49.4) | 7.0 (50.6) | 4.4 (57.0) |
| <I/σ(I)> | 24.5 (3.8) | 26.0 (2.8) | 20.2 (2.7) | 32.8 (2.8) |
| Redundancy | 8.5 (4.9) | 7.2 (5.2) | 8.2 (5.0) | 5.0 (4.9) |

| Sequences | dICS1 | dICS2 | RNAsub | DNAsub |
| --- | --- | --- | --- | --- |
| PDB code | 9OL0 | 9OL1 | 9OL2 | 9OL3 |
| Beamline | NSLS-II  17-ID-2 | NSLS-II  17-ID-2 | APS 23-ID-B | APS 23-ID-B |
| Wavelength (Å) | 0.97933 | 0.97933 | 1.033175 | 1.033175 |
| Space group | H32 | H32 | P321 | P321 |
| Unit cell parameters (Å, °) | 41.3, 41.3, 124.4,  90, 90, 120 | 40.9, 40.9, 125.0,  90, 90, 120 | 49.6, 49.6, 81.9,  90, 90, 120 | 48.2, 48.2, 82.7,  90, 90, 120 |
| Resolution range (Å) | 34.4-1.15 (1.17-1.15) | 34.1-1.24 (1.26-1.24) | 50-1.53 (1.56-1.53) | 50-1.56 (1.59-1.56) |
| Unique reflections | 14617 (819) | 11727 (572) | 17783 (757) | 15265 (433) |
| Completeness (%) | 98.1 (94.5) | 100 (100) | 96.9 (84.9) | 92.4 (55.2) |
| R_merge_ (%) | 6.1 (126) | 12.8 (200) | 7.9 (41.5) | 5.2 (43.8) |
| <I/σ(I)> | 14.9 (1.5) | 12.5 (2.0) | 15.9 (2.8) | 49.6 (2.6) |
| Redundancy | 9.8 (8.2) | 17.6 (14.7) | 5.9 (4.8) | 6.5 (3.7) |

**Table S4.** Data Refinement Statistics

| Sequences | dAU1 | dAU2 | dAUS1 | dAUS2 |
| --- | --- | --- | --- | --- |
| PDB code | 9OKS | 9OKT | 9OKU | 9OKV |
| RNA strands per asymmetric unit | 1 | 1 | 1 | 1 |
| Resolution range (Å) | 41.86-1.50 | 34.13-1.32 | 41.52-1.48 | 41.46-1.23 |
| Number of reflections | 6402 | 8918 | 6761 | 11009 |
| R_work_ (%) | 16.6 | 18.3 | 17.0 | 18.3 |
| R_free_ (%) | 21.0 | 20.6 | 21.1 | 22.7 |
| Bond length R.M.S. (Å) | 0.010 | 0.010 | 0.011 | 0.013 |
| Bond angle R.M.S. (°) | 2.01 | 2.30 | 1.98 | 2.04 |
| Average B-factors (Å^2^) | 10.3 | 11.9 | 11.5 | 12.1 |

| Sequences | dGC1 | dGC2 | dGCS1 | dGCS2 |
| --- | --- | --- | --- | --- |
| PDB code | 9OKW | 9OKX | 9OKY | 9OKZ |
| RNA strands per asymmetric unit | 1 | 1 | 2 | 6 |
| Resolution range (Å) | 41.53-1.54 | 41.54-1.38 | 30.33-1.48 | 36.81-1.92 |
| Number of reflections |  | 7991 | 14221 | 16740 |
| R_work_ (%) | 19.9 | 17.7 | 22.0 | 18.7 |
| R_free_ (%) | 24.8 | 21.5 | 24.8 | 25.7 |
| Bond length R.M.S. (Å) | 0.002 | 0.008 | 0.012 | 0.007 |
| Bond angle R.M.S. (°) | 0.48 | 1.86 | 2.21 | 2.06 |
| Average B-factors (Å^2^) | 14.8 | 10.6 | 12.2 | 14.9 |

| Sequences | dICS1 | dICS2 | RNAsub | DNAsub |
| --- | --- | --- | --- | --- |
| PDB code | 9OL0 | 9OL1 | 9OL2 | 9OL3 |
| RNA strands per asymmetric unit | 1 | 1 | 2 | 2 |
| Resolution range (Å) | 34.37-1.15 | 34.08-1.24 | 42.95-1.53 | 41.78-1.56 |
| Number of reflections | 14594 | 10551 | 16762 | 14497 |
| R_work_ (%) | 20.3 | 18.6 | 19.8 | 20.4 |
| R_free_ (%) | 23.3 | 22.9 | 23.4 | 24.9 |
| Bond length R.M.S. (Å) | 0.013 | 0.012 | 0.020 | 0.020 |
| Bond angle R.M.S. (°) | 1.77 | 2.08 | 3.04 | 2.93 |
| Average B-factors (Å^2^) | 20.0 | 12.6 | 15.9 | 48.1 |

**Table S5. Combination of the primer, template, blocker and complementary oligonucleotides used in the Michaelis-Menten analysis of primer extension reactions.**

| Bridged dinucleotide | Template Sequence | Primer | Template | Blocker | Complementary DNA |
| --- | --- | --- | --- | --- | --- |
| s^2^U*s^2^U | -AA- | DL-30 | LA-121 | LA-111 | dCLA-121 |
| s^2^U*s^2^U | -dAdA- | DL-30 | dAT26 | LA-111 | dCLA-121 |
| s^2^C*s^2^C | -GG- | DL-30 | LA-124 | LA-111 | dCLA-124 |
| s^2^C*s^2^C | -dGdG- | DL-30 | dGT26 | LA-111 | dCLA-124 |
| s^2^C*s^2^C | -II- | DL-30 | IIT26 | LA-111 | dCLA-123 |
| s^2^C*s^2^C | -dIdI- | DL-30 | dIT26 | LA-111 | dCLA-123 |
| A*A | -s^2^Us^2^U- | DL-30 | S2UT | LA-111 | dCLA-122 |
| dA*dA | -s^2^Us^2^U- | DL-30 | S2UT | LA-111 | dCLA-122 |
| G*G | -s^2^Cs^2^C- | DL-30 | S2CT | LA-111 | dCLA-123 |
| dG*dG | -s^2^Cs^2^C- | DL-30 | S2CT | LA-111 | dCLA-123 |
| I*I | -s^2^Cs^2^C- | DL-30 | S2CT | LA-111 | dCLA-123 |
| dI*dI | -s^2^Cs^2^C- | DL-30 | S2CT | LA-111 | dCLA-123 |

**Table S6. Sequences of oligonucleotides used in the Michaelis-Menten analysis of primer extension reactions.**

| Name | Role | Source | Type | Sequence (5ʹ→ 3ʹ) |
| --- | --- | --- | --- | --- |
| DL-30 | Primer | IDT | RNA | /FAM/AGU GAG UAA CGG |
| LA-111 | Blocker | IDT | RNA | G AUG UCA GAU AU |
| IIT26 | Template | In-house | RNA | AU AUC UGA CAU C**II** CCG UUA CUC ACU |
| dIT26 | Template | In-house | RNA | AU AUC UGA CAU C**dI****dI** CCG UUA CUC ACU |
| LA-121 | Template | IDT | RNA | AU AUC UGA CAU C**AA** CCG UUA CUC ACU |
| dAT26 | Template | In-house | RNA | AU AUC UGA CAU C**dAdA** CCG UUA CUC ACU |
| dCLA-121 | Complementary strand | IDT | DNA | AGT GAG TAA CGG **TT**G ATG TCA GAT AT |
| LA-122 | Template | IDT | RNA | AU AUC UGA CAU C**UU** CCG UUA CUC ACU |
| dCLA-122 | Complementary strand | IDT | DNA | AGT GAG TAA CGG **AA**G ATG TCA GAT AT |
| LA-123 | Template | IDT | RNA | AU AUC UGA CAU C**CC** CCG UUA CUC ACU |
| dCLA-123 | Complementary strand | IDT | DNA | AGT GAG TAA CGG **GG**G ATG TCA GAT AT |
| LA-124 | Template | IDT | RNA | AU AUC UGA CAU C**GG** CCG UUA CUC ACU |
| dGT26 | Template | In-house | RNA | AU AUC UGA CAU C**dGdG** CCG UUA CUC ACU |
| dCLA-124 | Complementarystrand | IDT | DNA | AGT GAG TAA CGG **CC**G ATG TCA GAT AT |
| 2SCT | Template | In-house | RNA | AU AUC UGA CAU C**s^2^Cs^2^C** CCG UUA CUC ACU |
| 2SUT | Template | In-house | RNA | AU AUC UGA CAU C**s^2^Us^2^U** CCG UUA CUC ACU |

**Reference**

1.	Ding, D., Zhou, L., Giurgiu, C. and Szostak, J.W. (2022) Kinetic explanations for the sequence biases observed in the nonenzymatic copying of RNA templates. *Nucleic Acids Research*, **50**, 35-45.

2.	Fang, Z., Jia, X., Xing, Y. and Szostak, J.W. (2025) Nonenzymatic RNA copying with a potentially primordial genetic alphabet. *Proceedings of the National Academy of Sciences*, **122**, e2505720122.
